# An agentic AI environment to support biomedical research in collaborative academic environments

**DOI:** 10.64898/2026.09.24.753565

**Authors:** Timothy Wu, Tyler S Browne, Mohamed Meawad, Hagar E Emam, Noor H Simsam, Yubing Xia, Amani Jaafar, Athena Ma, Bushra Kabbani, Vaibhav Gupta, Eliseos J Mucaki, Stephanie L Bishop, David R Edgell, Greg Gloor, Kun Ping Lu, Lynn Weir, Xiao Zhen Zhou, Vanessa Dumeaux, Michael T Hallett

**Affiliations:** Department of Biochemistry, Western University, 1151 Richmond St., London, N6A 3K7, Ontario, Canada; Department of Anatomy and Cell Biology, Western University, 1151 Richmond St., London, N6A 3K7, Ontario, Canada; Department of Oncology, Western University, 1151 Richmond St., London, N6A 3K7, Ontario, Canada; Robarts Research Institute, Schulich School of Medicine & Dentistry, Western University, London, N6G 2V4, Ontario, Canada; Department of Pathology and Laboratory Medicine, Schulich School of Medicine & Dentistry, Western University, London, N6A 5C1, Ontario, Canada; Lawson Health Research Institute, Western University, London, N6C 2R5, Ontario, Canada

**Keywords:** agentic AI, biomedical research, agentic operating system, security, traceability, collaborative research environment

## Abstract

Agentic artificial intelligence (AAI) is increasingly used by biomedical scientists, where it has substantially lowered the difficulty of integrating computational, statistical and data-science approaches into day-to-day research activities, assisting with project design, analysis and manuscript preparation. Significant barriers remain, however, which are unlikely to be removed solely by releasing more capable AI models. Meaningful use still demands a technical fluency that many biomedical researchers lack: a PI and their lab members must learn how to use the AAI system, establish standard operating procedures (SOPs) for managing, sharing and curating data, and enforce rules that address data privacy and security. Existing commercial AAI systems, which were designed primarily for software engineering purposes, need to be significantly re-purposed for biomedical-related data science.

We present Murmurent, shared software that sits beneath the agentic AI and provides the skills, procedures and protections to address the concerns above, removing the need for a lab to develop them independently. This includes (1) infrastructure to support projects and so-called *choreographies* involving multiple group members; (2) a set of *specialized agents* each dedicated to typical biomedical data science tasks including software and statistical development, data visualization, literature search, equity, diversity, inclusion, and decolonization (EDID) review, and others; (3) a tiered *memory* which can retain important research decisions, intermediate, derived outputs and sensitive data, and which can exploit this information to automatically build better contexts in AAI sessions; (4) *traceability* records which are used by the AAI session to better plan and execute data analyses and software builds by, for example, helping to avoid repeating decisions that lead to dead-ends; (5) *enforcement* of SOPs for data maintenance and data governance across all lab members; and (6) *multi-user capacity* to allow groups to interact and collaborate.

We use the system to identify putative inhibitors of Peptidyl-prolyl cis–trans Isomerase NIMA-interacting 1 (Pin1), a protein with a shallow catalytic site making it difficult to target. We describe the construction of several approaches to identify Pin1 inhibitors and the results they yield.

Murmurent is open source.

## 1 Introduction

Agentic artificial intelligence (AAI) has become a routine instrument in biomedical research, contributing to how experiments are scheduled and executed in high-throughput laboratories, how the resulting data are analysed and communicated, and how candidates are prioritized in areas such as drug discovery [1]. Three families of AI-augmented research dominate the published literature.

First, *lab-in-the-loop* workflows place a statistical or machine-learning predictor in an iterative design– make–test–analyse (DMTA) cycle. Using biotherapeutic discovery as an example, the model proposes candidate molecules or conditions, a wet lab tests them, and the predictor is updated on the feedback. Lab-in-the-loop drove much of the recent success of machine learning in biotherapeutics, where its most recent incarnations replace the classical predictor with a large language model (LLM) that plans the next experiment, queries chemistry and bioactivity tools, and drafts protocols rather than only emitting ranked candidates [2–4].

Second, *self-driving* or closed-loop laboratories take the DMTA loop one step further by replacing the human-operated wet lab with robotic execution, closing the hypothesis–experiment loop entirely in hardware [5–7]. Such systems are expected to have substantial impact in synthetic biology [8], in materials-science combinatorial discovery [9], and in the governance of automated experimentation in general [10].

Third, *multi-agent scientific-reasoning systems* chain together literature retrieval, code execution, and writing without robotic actuation. These systems are the clearest examples of agentic AI in biomedicine: LLMs perform actions on a researcher’s behalf by reading and writing files, making decisions, and proposing multi-step plans. For example, *Collaborative LLM-Agent Drug Discovery* (CLADD, Genentech Inc.) deploys specialized LLM agents backed by retrieval-augmented generation (RAG) [11] to integrate domain-specific knowledge into drug–target interaction prediction [12]. The AI Scientist-v2 from Sakana AI generates ideas, writes code, and produces manuscripts for a fixed class of machine-learning experiments [13, 14]; ChemCrow augments an LLM with 18 chemistry tools [15]. PaperQA2 reaches superhuman accuracy on literature question-answering [16]; its successor framework Aviary generalizes the agent to a gymnasium of scientific-task environments including DNA cloning and protein-stability prediction [17]. STORM curates structured reports through simulated expert interviews [18] and frameworks such as AutoGen [19] supply the agent-coordination scaffolding on which many of these systems run. Hybrid systems that combine multi-agent reasoning with a wet-lab validation step, for example, Robin [20] and AI co-scientist (Google DeepMind) [21], close the hypothesis-to-validation loop within a single workflow, reducing the rate of hallucinations and generating candidates which have been validated experimentally.

The three families of approaches share several limitations. To date, each system tends to be scoped to a single, well-defined end-point e.g. discovery of a biotherapeutic for a specific disease [22–26]. This is perhaps not surprising given that the paradigms often grew out of industrial pipelines, which tend to be “linear” in nature: a fixed funnel (target identification, lead discovery, lead optimization, preclinical testing) has a single success criterion, and AI is inserted as an accelerant at one or more stages. The linearization persists even in systems described as academic; Robin, for example, is vertically integrated around a single disease end-point within one team’s infrastructure, with no mechanism for sharing agents, data invariants, or governance across independent users and laboratories. Academic environments, however, tend to be heterogeneous federations of laboratories and core facilities (e.g. genomics, proteomics, imaging, biostatistics) that form dynamic collaborations around shifting scientific questions, each with its own data, success criteria, and governance. The user populations are equally heterogeneous, ranging from undergraduate trainees to research associates, with widely varying programming experience and limited information-technology or bioinformatics background. Few academic centres have a bioinformatics core large enough to serve every group’s analytical needs on demand. Group *sovereignty* is a defining feature: each lab owns its data, sets its own success criteria, and answers to its own granting agencies. This combination of heterogeneity and autonomy is incompatible with governance approaches that route everyone’s work through a single central controller (a model often called *orchestration*), since no such controller can understand every project well enough to micro-manage it [27].

In academic environments, security, privacy, and intellectual property are acute concerns: proprietary molecules, unpublished results, and patient-derived data cannot simply be sent to a public LLM, because inadvertent memorization of training-time inputs has been demonstrated to occur [28]. Hospital data-use agreements strictly forbid disclosure to third-party processors and public submission can constitute prior art that defeats patentability. Academic centres also have a mandate to develop and disseminate new computational methods with a vested interest in integrating externally published tools into their software pipelines quickly. A workable centre-scale infrastructure must therefore interface cleanly with the commercial systems researchers already use (e.g. communication platforms, code repositories, inventory management software) and with internal institutional systems (e.g. financial reporting, core-facility billing, lab information-management systems), rather than displace them. Finally, health inequality [29], equity, diversity, inclusion, and decolonization (EDID) review, together with sex- and gender-based analysis plus (SGBA+) [30], are explicit requirements of major research funders and peer-reviewed venues across multiple jurisdictions; an academic-grade agentic system must integrate this review as a first-class activity rather than as an afterthought.

We introduce *Murmurent*, shared agentic AI infrastructure designed specifically to support molecular biomedical research in an academic collaborative environment. Murmurent aims to direct the power of agentic AI towards making agentic AI itself more accessible to biomedical researchers regardless of stage of training or lack of background in quantitative methods. It is best understood as an *agentic operating system*; that is, the layer beneath agentic AI, which provides the identities, permissions, shared memory, and data-governance invariants that allow many such agents to coexist safely on shared infrastructure. Murmurent implements an alternative to the orchestration model described above; our *choreography* model provides the capacity for each group to build its own approaches to a given problem using shared agents and rules. We first present the design principles of Murmurent, highlighting key properties of the system that extend beyond default AAI command line interfaces (CLIs). A worked example related to biotherapeutic exploration is introduced at the outset of the presentation to help motivate each property to non-experts and to show how they can independently create such bioinformatic systems for their target problems.

## 2 Results

### 2.1 Overview of the agentic AI operating system

Murmurent is implemented as a layer above a commercially available agentic AI system (Claude Code, CC; Anthropic) and extends it along six dimensions. Figure 1 provides an overview of its general structure.

**Fig. 1.**
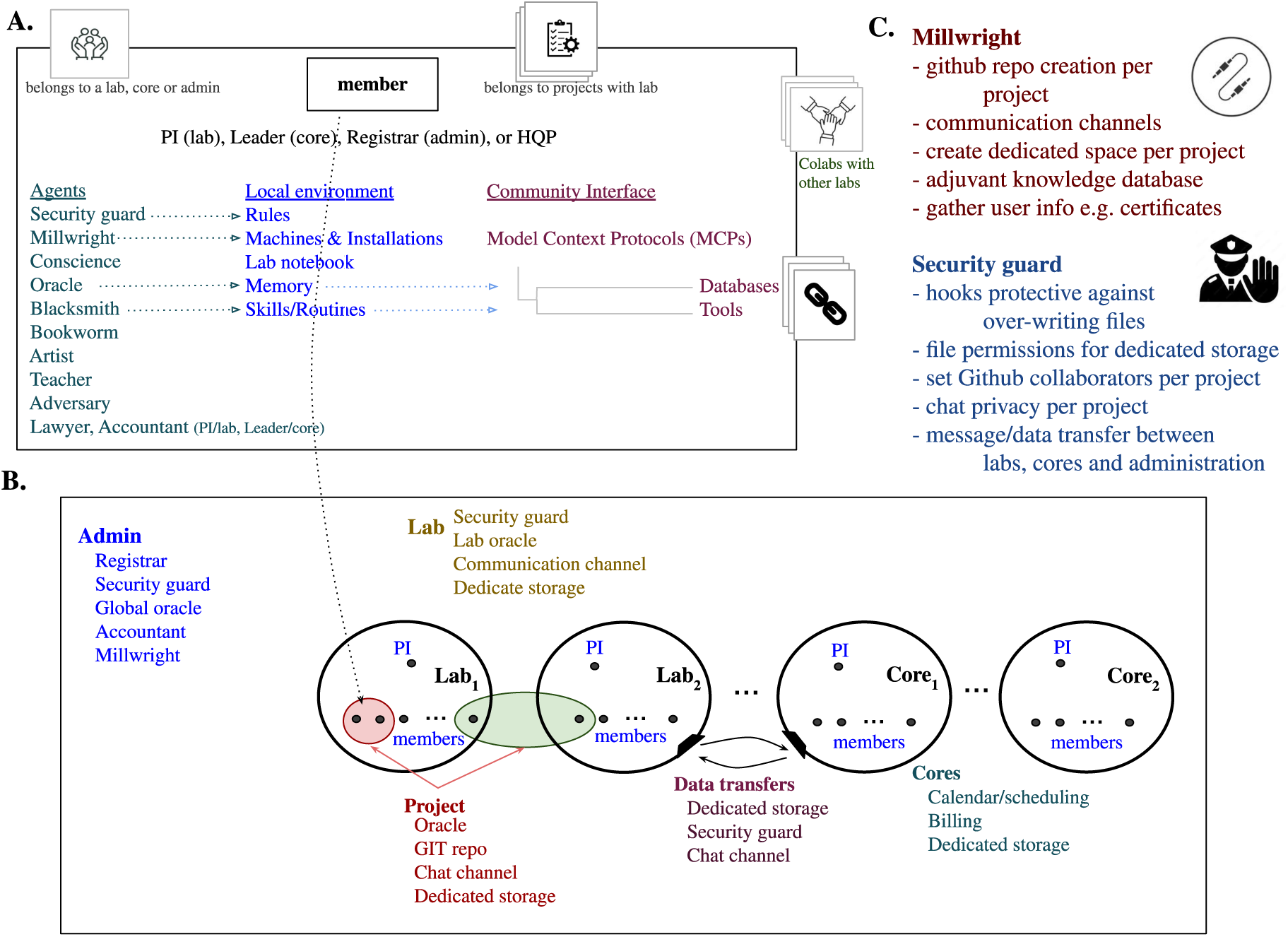
Architecture of Murmurent. **(A) Member view.** Every Murmurent user (PI of a group (lab or core), group members, mayor in administration) draws on the same resource families: reference *agents* for common day-to-day tasks which can be easily customized for the user’s domain-specific challenges, rules for security/privacy, secure machines and installations, an electronic lab notebook, different types of memory, along with pre-developed skills/routines/workflows available in most agentic AI systems; and a community interface of Model Context Protocol (MCP) servers that expose external databases and institutional services into a shared choreography. **(B) Social structure.** An administration tier provides oversight across the labs and cores; each such group carries its own Security Guard, Oracle, chat channel, and dedicated storage. Projects can be scoped to a single lab (red) or span multiple labs (green). **(C) Detail on two foundational agents.** The *Millwright* handles provisioning (per-project GitHub repositories and chat channels, dedicated storage, adjuvant knowledge databases). The *Security Guard* enforces the data-governance invariants described in Methods (hooks against overwriting files under `immutable/` or `append only/`, storage permissions, GitHub collaborator scope, chat privacy, inter-entity message and data transfer auditing).

#### 1. Projects and choreographies

Work (e.g. data analysis, software, literature search, manuscript development) is organized into projects. A project provides a group of users with shared dedicated communications channels, persistent storage and a written record of all the performed work. When a project is initiated in Murmurent, it creates a private chat channel (Slack), GitHub repository, named directories on user-shared drives to provide private, secure storage space for large-scale data, and documents generated by CC that track design decisions and the state of software builds and long-running computations. A cryptographic certificate is issued, which allows Murmurent to maintain control over which users have access to the project’s resources. Each project also carries a *charter*, a short plain-text file at the root of its repository that records the project’s name, its sensitivity tier, its lead, and a paragraph stating its purpose together with any standing caveat about the data. A second file is used to indicate that the folder is governed by Murmurent, and therefore is subject to specific restrictions and procedures e.g. screening for patient identifiers (§2.6). A hook injects the charter’s opening paragraph, the project’s name and sensitivity, the reader’s role in the project (§2.4, §4.4), and the rules that screen for protected and sensitive information (§2.6).

Murmurent also supports *choreographies*. A choreography begins with a significant question or challenge whose solution likely requires a complex, multi-component approach. The example below asks for inhibitors of Pin1, which could be approached in different ways e.g. by pursuing covalent or non-covalent chemistry, by generating candidates from scratch with a deep-learning model, or by modifying a known inhibitor (§2.2). Each user may answer the challenge of the choreography with an approach of their choosing. Murmurent facilitates interactions between the approaches and decides the terms of comparison: which quantities are reported, in what units, how values are interpreted, how uncertainty is expressed, and what serves as a control. The Pin1 choreography settled its control molecules, decoys and tests of significance before any of its four approaches were built. Three things follow from distributing the work this way: (i) Each group maintains authority over its own data and its own methods while still answering a shared question; (ii) It motivates finding consensus on issues such as measures of goodness, controls and other items required for integrative analyses; and (iii) It allows incommensurable quantities to coexist and recognizes that such differences may themselves be informative. In essence, it is the question which brings the users together without a single controller.

#### 2. Specialized agents

Every Murmurent user inherits the same versioned, centrally reviewed set of reference agents, each designed for one aspect of biomedical and basic bioscience research (§2.3). This saves the user from building and evolving their own agents. It also allows a PI to distribute the common agents, and subsequent improvements to the agents, easily across all lab members. Research-specific agents cover statistical, computational and data-science methodology; visualization, communication and training; literature retrieval, biomedical-database integration and teaching (§A.1); and the curation of personal and group institutional memory. A second set performs independent, adversarial review of each step of the analysis generated by the system. This includes intellectual-property and freedom-to-operate assessment, audit of secrets, protected health information file permissions (§2.6), and EDID/SGBA+ review (§A.2). A third set maintains the infrastructure the first two run on: provisioning and periodic verification of accounts, repositories, storage and communication channels (§2.7).

#### 3. Memory

Murmurent separates memory into three tiers, each stored, used and governed differently (§2.4). *Tier 1, contextual memory*, is what an agent can see during one session, and it is discarded when the session ends. Murmurent fills it at the start of every session with the project’s history, the design decisions already taken, and the rules the project works under. This frees the researcher from restating project conventions in each new session, minimizes the need for agents to guess how they should behave and helps to avoid repeated mistakes and wasted effort during future builds. *Tier 2, persistent memory*, holds what the work has learned, including observations, decisions, negative results and methodological choices, as markdown files in the researcher’s own account, available to later sessions and to other projects, and written through the Oracle agent. *Tier 3, external memory*, holds bulk data (e.g. raw genomic files, large-scale analyses, visualizations and subsequent reports) on the filesystem under conventions which follow a lab’s declared standard operating procedures (SOPs) and make critical files difficult to overwrite or delete.

#### 4. Traceability

Every tool call, design decision, significant failure and administrative action is recorded, which allows Murmurent to assemble a context that steers a later session away from a dead end the project has already been down (§2.5).

#### 5. Enforcement

Data-governance rules are applied when an agent acts rather than asked of it. Each tool call is inspected before it runs and refused if it would break a rule, whatever the model reasoned, however the request was phrased, and whether or not the member knows the rule exists (§2.6).

#### 6. Multi-user operation

Data, results and decision records can be shared between members. Murmurent supports groups led by a principal investigator (PI), projects that form within or across groups, and a centre that houses several groups. It creates the secure communication channel, code repository and protected storage to facilitate each of these needs (§2.7).

The subsections that follow discuss each of these dimensions but first we introduce an example biotherapeutics choreography in order to better motivate each property.

### 2.2 An example Murmurent choreography: *Pin1* inhibitor

The central challenge of our choreography is to find a small molecule that inhibits human Peptidyl-prolyl *cis*– *trans* Isomerase NIMA-interacting 1 (Pin1). Pin1 uniquely isomerizes phosphorylated Ser/Thr–Pro peptide bonds and has established roles in cancer and neurodegenerative disease, which makes it an attractive drug target [31]. It is however difficult to target for structural reasons: the catalytic site is a shallow, solvent-exposed groove rather than an enclosed pocket, so a bound ligand buries little surface area, and docking scores therefore separate binders from non-binders only weakly [32]. The little affinity that exists is concentrated in a small, strongly basic phosphate-binding subsite. That subsite rewards acidic phosphate mimetics, chemical groups that imitate the substrate’s phosphate. Such groups carry a negative charge at physiological pH, and charged molecules cross cell membranes poorly, so the same chemistry that buys affinity for an intracellular target is what keeps the compound out of the cell [33]. The site is also conformationally plastic, adapting to whatever occupies it [34, 35], so any single prepared structure is one state of the pocket rather than the pocket in general. Cys113 is what promises tractability: a covalent warhead could convert weak recognition into durable engagement [36].

A candidate inhibitor should ideally pass structural-alert and drug-likeness filtering, be synthetically accessible, and have a plausible, reproducible predicted binding mode rather than merely a favourable docking score. A covalent candidate should additionally engage Cys113 through a chemically valid mechanism whose electrophilic reactivity falls within a precedented range. These conditions are tracked per candidate rather than imposed as universal filters. In two of the approaches described below, two reported inhibitors are considered: sulfopin, a covalent chemical probe that engages the catalytic cysteine Cys113 [37], and all-*trans* retinoic acid (ATRA), a weak non-covalent binder [33]. Throughout our example choreography, docking makes use of one prepared Pin1 receptor (PDB 6VAJ), and every quantity we report for this choreography was measured on that receptor.

Four distinct approaches spanning the space from seed-free generation to expert-constrained covalent design are presented (Figure 2):

1. *T*_1_**, *de novo* structure-based generation.** A pocket-conditioned deep generative model (DiffSBDD) reads the Pin1 catalytic site and samples novel non-covalent ligands from scratch with no seed molecule [38]. The approach demonstrates bringing a complex, public deep-learning method under the choreography’s control, which includes released weights, a GPU environment, and docking-based rescoring, which a non-expert could not realistically get running alone.
2. *T*_2_**, derivative neighbourhood of ATRA.** Starting from ATRA, the approach (CReM) enumerates a degree-bounded neighbourhood of graph edits [39] and labels each non-covalent derivative through a cost-ordered ladder of cheap-to-expensive attributes, reducing the set to a shortlist through an internal deliberation loop.
3. *T*_3_**, single-warhead decoration of sulfopin.** Holding sulfopin’s scaffold and one expert-chosen covalent warhead fixed as hard constraints, a generative model (REINVENT) [40] decorates only the remaining R-group position. This is the most constrained approach: one degree of freedom, with the covalent mechanism fixed before the generator runs.
4. *T*_4_**, combinatorial warhead** × **R-group search.** Holding only the sulfopin core fixed, the approach enumerates a covalent warhead library against an R-group library, then filters, docks covalently at Cys113, and triages each warhead class against a reactivity window bounded by known covalent Pin1 actives. This is the fullest expert-plus-tooling covalent approach, and the only one addressing every condition above.

**Fig. 2.**
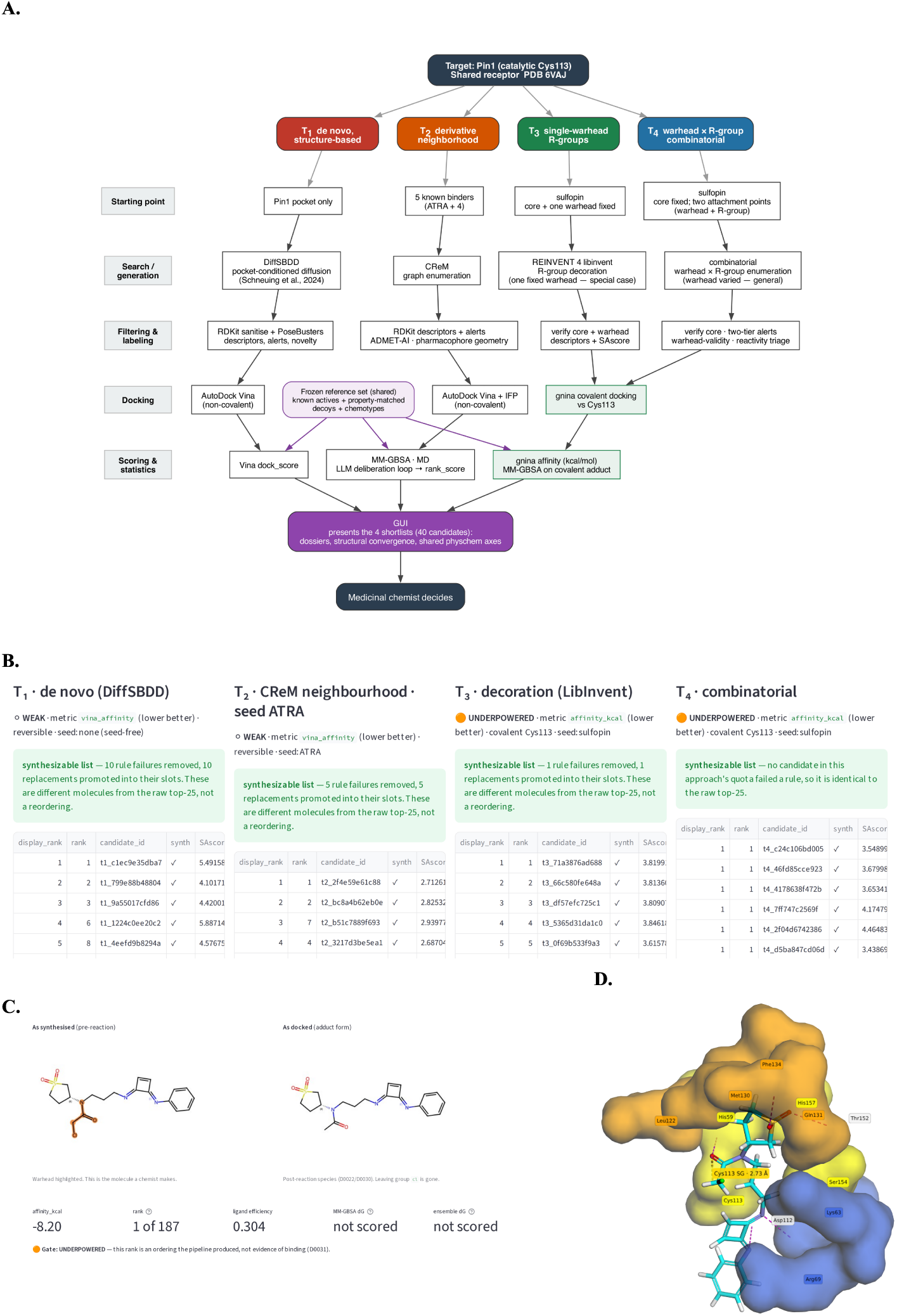
The four *Pin1* approaches. **A.** All four approaches start from one shared target, the Pin1 catalytic Cys113 on a single prepared receptor structure (PDB 6VAJ), and successively address five goals (left, gray text). Each approach (column) differs in how much expert constraint it carries, from *`T`*_1_, which is given only the pocket, to *T*_4_, which holds the sulfopin core fixed and varies both a covalent warhead and an R-group. Two things are deliberately shared rather than duplicated per column: the frozen reference set of known actives, property-matched decoys and chemotypes (centre), which every scoring stage is measured against, and the descriptor and alert code invoked in the filtering row. The three scoring stages remain distinct quantities and are not merged into a single ranking. They meet only at the interface. **B.** In a GUI generated by the Artist, the four shortlists are depicted side by side, with the properties each candidate carries and the verdict its control issued. **C.** One candidate’s structure, as synthesized and as docked. **D.** That candidate’s predicted pose in the Pin1 active site, rendered with py3Dmol, integrated by the Artist into the GUI. Residues within 4.5 Å of the ligand are labelled; dashed lines are polar contacts, drawn where both partners are nitrogen or oxygen and within 3.6 Å, and are measured. The gold dashed line reports the closest approach of the ligand to the catalytic Cys113 sulfur; it is a measured distance, not a modelled bond. The molecule in **C** and **D** is *T*_4_’s first-ranked candidate under the size-decorrelated metric the shortlist is ordered by, reproduced to illustrate the interface.

All four approaches are judged against one shared control. The control is a set of decoy molecules chosen to resemble the known Pin1 binders in size, greasiness and charge, the properties a docking score responds to most strongly [41, 42]. An approach that claims to find binders should rank the known binders above these decoys.

#### How the choreography proceeded

Four people from different labs developed the approaches in independent sessions of CC. Each championed a single approach, deciding upon the methods, models and software with iterative feedback provided by CC and Murmurent at each step. No centralized agent judged one approach superior to another. This is what distinguishes a choreography from a centralized pipeline with a conductor. The approaches share only a common goal and Murmurent, which provides constant assistance for building and executing each approach and which attempts to integrate the approaches when it makes sense.

Murmurent started by deciding on shared standards across the four approaches including the use of conventions (e.g. for representing molecules in SMILES nomenclature, for adding hydrogens and for attaching a molecule to Cys113), measurements (e.g. physical characteristics, synthesizability) and methods for comparison (e.g. level of evidence required to claim a candidate as statistically significant). The four approaches were instructed to use a set of published Pin1 binders with distinct chemotypes as a control. This was assembled by the Bookworm through database and literature search. Murmurent suggested that the four approaches use the same prepared structure of the Pin1 protein, with a search region sized to each mechanism: a tighter box for the covalent approaches compared to non-covalent approaches with the understanding that scores computed in different volumes are not directly comparable.

Each such decision was written down in the *traceability* infrastructure provided by Murmurent (§2.5). Output analyses are shared between the approaches through Murmurent’s memory model (Tier 2, §2.4), and all development/computations were performed under uniform security and privacy protocols (§2.6). If the four approaches had been developed separately without Murmurent, each would have arrived with its own reasonable but private definition of novelty, challenging the feasibility of downstream comparisons.

Murmurent provided several additional forms of support during execution of the code. First, because all four used the same code for the shared steps, any coding errors were exposed and corrected for everyone. For example, a defect in the shared module that flags problematic chemical groups was noticed only because it induced an extreme percentage (97.7%) of candidates being rejected for one of the four approaches. Repairing it corrected every approach at once. This was a semantic bug that may have been very difficult to find if the approaches had been developed separately. Second, the Adversary assessed the feasibility and the resource demands of each proposed computation, and kept the long calculations running across several days and several iterations (§2.3).

The four sets of results are presented side by side along with supporting evidence of their candidacy in a unified graphical user interface built by the Artist (Figure 2 B, C, D). It is the user who weighs the evidence and picks the compounds worth expensive lab work. If a new, fifth approach were to be implemented, it would start with the decision records, the reference set of published binders, and the controls rather than starting from scratch.

We describe how Murmurent assists with making final conclusions below (§2.8) after the properties of Murmurent are described in more detail.

### 2.3 Murmurent provides a robust set of AI agents

Every agent closes its work with a *verdict*: a single categorical judgement drawn from a short list fixed in advance for that agent, placed in the first line of its reply and before any supporting detail. The Security Guard answers *Clear*, *Concerns* or *Blocked*; the Adversary *Pass*, *Questions* or *Reject*; the Teacher *Explained* or *Gap*. The lists are deliberately short, and in some cases lack a middle term, so that an agent which cannot support a favourable answer has to return an unfavourable one rather than qualify its way between the two. Three things follow. A researcher reads the conclusion without reading the report. A later stage of a build can act on the judgement without interpreting prose, which is how one approach in a choreography learns that another has failed its controls. And because the same vocabulary is used by every member of the centre, two reviews of the same analysis are comparable even when different people requested them (§4.2).

We briefly summarize the role of each agent provided by default in the Murmurent environment, but stress that these agents can be easily tailored to any specific environment or project, and new agents are easily added to the system.

Several agents are dedicated to various aspects of research and production.

The <u>Blacksmith</u> is the workhorse that focuses on the analysis of data and development of software. It should be tailored to the needs of each lab with domain-specific tools and resources. It should also be set up to follow lab protocols and preferences including, for example, a specific programming language, statistical approaches and user interface conventions. Because it usually writes the build plan, it is assigned the most capable model available. Its definition also states what it may not do: figures go to the Artist, and cross-referencing a prediction against published work goes to the Bookworm, so each task reaches the right agent without the researcher directing traffic. In the Pin1 choreography the Blacksmith carried the bulk of the computation, docking and ranking candidate molecules across the four approaches, and monitoring the computations across 5 GPUs over the course of several weeks.

The <u>Bookworm</u> focuses on literature survey and external, publicly available data. Because the Bookworm’s primary role is to interact with databases such as PubMed, CrossRef and preprint servers, it is one of the few agents granted web access Most agents are not allowed web access in order to minimize the chance that intellectual property is accidentally transferred to a third party. Every claim it returns must carry the database and the accession or PubMed identifier it came from, and it is required to declare any conflict it meets between sources rather than silently prefer one. References are added to the lab’s reference manager, by default Zotero. In the Pin1 choreography the Bookworm assembled the reference set of published Pin1 binders on which the shared control is built, together with the same-chemotype molecule pools the control uses as negatives.

The <u>Artist</u> is dedicated to representing data through statistical plots, academic presentations, animations and educational videos. The group-wise shared aspect of the Artist facilitates more organized, systematic communication of results and can provide a common “look and feel” (e.g. group or institutional colour schemes and logos) although it too may be progressively updated by lab members with new techniques as they arise. The definition of the agent specifies the plotting library, default figure dimensions, colour map, presentation format, and good examples from third parties the user wishes the agent to consider. In the Pin1 choreography the Artist built the interface through which the four shortlists are presented side by side. It shows each candidate’s predicted binding mode, flags molecules found independently by more than one approach and embeds well-established molecular drawing tools.

The <u>Teacher</u> has three modes. (i) It can *debrief* the user when an agent produces an over-complicated or a jargon-filled answer to a prompt. (ii) The Teacher can also be asked to *explain* a method, paper, statistical idea, codebase or design decision for a technically competent but adjacent-field audience, optionally rendering a self-contained annotatable page with a self-grading quiz. (iii) It can design a multi-sectioned *course* for the user to assist them with more long-term learning of a specific concept, field or framework. In this mode, it sources its reading through the Bookworm rather than the open web, and keeps a per-learner record so that material a member has already demonstrated is not taught again. All three modes are illustrated in Figure S1 and § A.1.

Several agents are dedicated to memory of important findings, events and decision making in a project.

The <u>Oracle and Lab Oracle</u> are the agents through which researchers write to and read from Tier 2 memory (§2.4). Together they hold what has been learned across all projects the user or lab has been a part of. The Oracle is personal, processing each member’s own record of findings, hypotheses, negative results and the reasoning behind experimental choices. The user tells the Oracle which such facts should be remembered. This information is stored as markdown in the user’s (Obsidian [43]) vault. The Lab Oracle is similar to the personal Oracle but shared across the group and read-only to agents; an entry reaches it only when a group member proposes it and the PI approves. Because both are plain text under version control, a PI can audit them (§2.4). Every entry carries mandatory frontmatter information (title, date, project, sensitivity, tags and sources) which can be used as filters when querying. Every entry also carries one of three sensitivity tiers. An entry marked `standard` may be promoted to the group. One marked `restricted` or `protected` may not, and the publication tool refuses it.

Several agents are dedicated to reviewing, adjudicating and optimizing the output from the other agents.

The <u>Adversary</u> is dedicated to critiquing the collective findings of the other agents, primarily the Blacksmith, Bookworm and Artist, in a dynamic, continuous manner. Multi-agent and adversarial-critique approaches which instigate debate between an ensemble of agents have been shown to reduce LLM hallucinations, improve factual validity and increase the depth of answers [44, 45]. The development of effective adversarial agentic approaches remains challenging [46] and we expect our Adversary to evolve significantly with experience. By default, the Adversary in Murmurent insists on certain technical/experimental checks e.g. it will examine the distinctiveness of the training and test sets in machine learning settings, and verify that assertions made by other agents are correct including, for example, literature references, calculations and assumptions. In the Pin1 choreography the Adversary rejected an intractable exhaustive enumeration and an unscoped docking loop before either consumed significant computational resources, and a later audit of the choreography’s own rescoring protocol found two defects in numbers the project was preparing to report (§2.8).

The <u>Conscience</u> periodically reviews experimental design, text and literature with a focus on EDID concerns and SGBA+ handling of sex, gender and other population variables (e.g. bias in experimental design, language, source selection, and presentation) in order to promote mindfulness across the members, identify limitations of either literature-derived work or future planned work, and to assist with the development of respectful communications (§A.2). In the default Murmurent, the Conscience is configured against the CIHR SGBA+ framework [30] and extensive provincial (Ontario) and federal (Canada) guidelines. The Conscience is wired to the Security Guard so that when, for example, a project involving sensitive data is blocked by Murmurent, the Conscience will supply an explanation as to why the operation was flagged as problematic.

The <u>Lawyer</u> specializes in freedom-to-operate searches, issues of patenting, governmental regulations for the secure storage of clinical and commercial data, and disclosure. The Lawyer provides a means to incorporate issues related to intellectual property and regulation into computational decision-making “on the fly” during workflow execution. Such continuous, ongoing input from the Lawyer can help, for example, to prioritize small molecules without patent issues. It searches patent collections in a fixed order: United States granted claim text is read first, then WIPO PatentScope, then Google Patents. The Canadian Patents Database, DEPATISnet and two secondary collections are examined only when a question names a specific jurisdiction. For each finding, the Lawyer reports how the status was established, and marks a status as inferred where it was derived rather than read from an authoritative record.

The <u>Security Guard</u> inspects what is about to leave a member’s control including, for example, code and data staged for a Git push, data objects prepared for a collaborator, and files whose permissions would expose them beyond their intended audience. It scans for credentials, for patterns matching protected health information, and for paths that governance rules place off limits (§2.6). Its patterns are set per deployment and per jurisdiction: alongside generic credential formats (API tokens, SSH and age keys, `.env`-style assignments, cloud key signatures), a project declared `protected` is additionally scanned for health identifiers. In the default Murmurent deployment, this corresponds to Ontario identifiers including numbers resembling an Ontario health card (OHIP), a hospital medical record number (MRN) or a social insurance number (SIN), and dates of birth adjacent to names. Private signing keys of users are treated as paths that may never appear in a Git diff, log or message under any circumstance. The Security Guard can also be configured to periodically check the entire complement of files for a user or group to ensure that permissions are set properly everywhere.

Finally, a subset of the agents focuses on building and maintaining infrastructure across the users, groups and administration.

The <u>Millwright</u> and the <u>Centre Millwright</u> are both dedicated to the installation and verification of all group and administrative protocols. The agents periodically and automatically perform background audits to identify whether infrastructure is improperly maintained including communication channels, Git-based repositories, project disk space, and project membership. Each group declares its own tools and storage layout (e.g. the GitHub CLI for repositories, the chat workspace in Slack for channels, and directory structures). Once the PI has decided on such lab protocols, the Millwright ensures as best possible that all members of the group follow this design. The Millwright greatly simplifies the installation and maintenance of data and software especially for non-experts. The Centre Millwright has a similar mandate but focuses on centre-wide protocols and procedures including the fair sharing of available computational resources.

The <u>Registrar</u> maintains a record of the groups (labs or cores) and their respective PIs across the centre. The Registrar is the authority on what that infrastructure is meant to reflect, and prompts the Centre Millwright to detect, report and correct deviations from this vision. The Registrar has a view across groups that no individual PI holds, which may have importance when the centre is required to track resource usage, plan resource extensions and distribute available computational resources (§2.7). Groups are opaque to the Registrar in order to maintain privacy. The records indicate only the membership of groups and projects and the Registrar is forbidden from peering inside any entity (notebooks, projects and personal records).

### 2.4 Memory within the agentic OS

In most agentic AI systems, memory is undifferentiated: the model’s prompt window holds the live conversation, and additional knowledge, when needed, is fetched on demand often facilitated by *RAG* systems, as in CLADD [12] and Robin [20], or by *ad hoc* tool calls in other systems. Either way, the result is a single stream of text presented to the model, with no structural distinction between live conversational state, the lab’s accumulated knowledge, and the bulk scientific data the analysis operates on. Murmurent instead separates memory into three explicit tiers, each with its own representation, access pattern, and governance.

#### Tier 1, contextual memory

This corresponds to the in-session working memory of the agent, consisting of the user’s prompt, the agent’s reasoning, and the immediate tool-call traces. Contextual memory is short-lived (within a single session) and fully delegated to the underlying agentic CLI. Murmurent makes two contributions at this tier. The first is session-level audit logging, through a hook that fires when a subagent finishes, producing the per-user activity log described in §2.6.

The second is that a session begins already knowing where it is by two different routes. A project’s `CLAUDE.md` file, which the agentic CLI reads at the start of every session, imports the project’s orientation document and its catalogue of previously diagnosed failure modes (§2.5) by path, so both are automatically present in full, every session. The decision records cannot be imported the same way in part due to their size. For example, the Pin1 choreography has 57 decision files, so importing them alone would fill the context. `CLAUDE.md` therefore carries three smaller things: the path to the directory, the format a record must follow, and one standing instruction for what to do on finding a defect. That instruction has three parts: repair the class of defect rather than the single instance, write a record that says why the wrong answer looked right, and add a guard capable of failing. Separately, a hook prepends the project’s name and sensitivity, the opening of its charter, and the member’s role in it. Together these approaches ensure that the reasoning survives in written form, so that repeated mistakes can be caught during future builds and analyses.

#### Tier 2, persistent memory

Durable, qualitative knowledge such as observations, decisions, near-miss results, literature summaries and methodological choices is recorded in a user-specific container, implemented as an Obsidian vault [43]: essentially a folder of files in markdown. The vault is structured as follows:

- `oracle/`, curated entries produced by the Oracle agent, each carrying mandatory frontmatter (title, date, project, sensitivity, tags, sources);
- `lab-notebook/`, dated entries with no frontmatter requirement, giving users a convenient means to track daily experiments with different types of information (text, PDFs, images, etc.);
- `murmurent data/`, arbitrary reference material (PDFs, spreadsheets, protocols, images etc.) that agents read on demand to condition their work;
- `maps-legends/`, the vault’s own taxonomy and index, which agents consult before writing so that tags and structure stay consistent across entries.

The personal and the group record are held separately, the first in the member’s own vault and the second in a repository the group owns and only the PI may write to. The split separates a potentially noisy stream of personal notes from the curated record the lab inherits across personnel turnover, and restricted entries are never promoted from the first to the second. The Oracle is the means by which knowledge enters the curated part of this tier. A researcher tells the Oracle what should be remembered, and the Oracle writes it as an entry with the required frontmatter, so that nothing reaches `oracle/` without a named author, a date and a sensitivity. The Lab Oracle is the corresponding read-only view of the group’s record, and `murmurent oracle publish` is the only path from the first to the second. Entries are queryable by project, date, tag, sensitivity and author through a small Model Context Protocol (MCP) server (§4.5), so any agent can retrieve them.

General-purpose LLMs have difficulty reliably retrieving from the open web, likely because real-world biomedical questions span heterogeneous data types and specialized knowledge sources. Previous efforts (CLADD [12] and Robin [20]) have addressed this with RAG [11]: a corpus is embedded into a vector index, and each query returns the text chunks nearest to it. Such a lookup is opaque: each call returns a few text chunks, with little indication of which document they came from, who wrote it, or under what sensitivity classification. It is also stateless: the vector index is a derived cache, regenerated whenever the corpus or embedding model changes, so it carries no history of what was known when, and a retrieval cannot be reproduced once the index has been rebuilt. Murmurent’s memory is instead plain markdown under version control, found by filtering on the frontmatter described above and keyword-scoring the body, so retrieval is reproducible by hand. A person can open the same file the agent read, see who recorded it and under what sensitivity, correct it in place, and recover the correction from the Git history.

The Pin1 choreography applies these conventions, with version control, to the shared files of the project, which are of three kinds. These files are not themselves Tier 2 memory: they live in the project’s repository rather than in a member’s vault, and they are written by the agents that produce them rather than through the Oracle. They follow Tier 2 conventions so that they carry the same provenance and can be audited in the same way. The first is the decision records described in §2.5. The second is the reference sets, such as the published Pin1 binders and the covalent warhead library. Each carries a version number, is never edited in place, and travels with a companion file naming its source, the agent that assembled it, and its known limitations. The third is a single file recording every external input the choreography depends on, each identified by a checksum of its contents, so a later run can confirm it used the same data.

All four approaches of the Pin1 choreography read a single common copy of the published binder set. At least two events occurred during development of the choreography that showed why Tier 2 conventions here pay off. The first event occurred during curation of the binder set itself where Murmurent found that two molecules had been taken from a database record for a different enzyme, threonine-tRNA ligase, in the belief that it was human Pin1. Both were removed. The two were non-covalent binders, and their removal left only three valid non-covalent actives. Because the set is assembled once and shared, one correction reached all four approaches simultaneously.

The second concerns a chemical structure the Bookworm could not confirm. *T*_4_ screens candidates against a reactivity window; that is, the range of reactivity measured for known covalent Pin1 binders (§2.2). Trust in that window is determined by the quality of the control molecules. One such molecule identified by the Bookworm appears only as a drawing in a supplementary figure of a manuscript with no matching entry in any public database. To include it in the analysis required reading the structure off the image “by eye”, a process that may introduce errors. To safeguard against this entry “polluting” downstream computations, the Bookworm recorded in the companion file that this entry was ‘unverified’ along with a short description as to why and how it could be resolved. Downstream code was designed to refuse to send an unverified row to the reactivity window. In this way, applying Tier 2 conventions to the companion file allowed questionable data to be treated specially and to provide the user with instruction as to how the situation could be resolved in order to gain an additional control molecule.

#### Tier 3, external memory

Common data files such as sequencing reads, mass-spectrometry runs, NMR FIDs, microscopy stacks and similar data objects are too large for the Oracle and too varied for a single index, so Murmurent stores such data under several filesystem conventions, each upheld for agents by a hook. For example, a read-only `immutable/`

`<project>/` tree is created for the original raw data obtained from instruments, collaborators, or public repositories, and an `append only/<project>/<experiment>/` tree for the outputs of analyses and experiments. Both are upheld for agents by CC hooks that refuse mutating tool calls on those paths (§A.3). Agents read bulk data with ordinary tools (e.g. `grep`, `cut`, `sort`, `head`). Where a store has a natural query interface, such as a relational schema for cohort metadata or a chemical-structure database, Murmurent wraps it as an MCP server rather than copying it into a prompt.

### 2.5 Traceability

The construction and execution of a choreography could take considerable time and involve multiple individuals. The reasoning behind each decision during this project is captured neither in the version-control history nor in the transcript of the conversation. This type of information is lost when sessions are aborted. However, a record of such information (e.g. alternatives considered, evidence for decision making, and on whose authority) would assist in future builds and avoid repeated mistakes. Murmurent responds to this challenge by constructing a series of files that outlive both the session and the person, in formats a human reads directly and a machine can validate. The Pin1 choreography instantiates this in three document kinds, which we describe here (see also §A.4 for examples).

#### Decision records

Each decision file (stored in decisions/) records a consequential choice made during the build or execution of a choreography. Murmurent judges which such decisions would be expensive or confusing to reverse, or difficult to explain months later, and ignores routine implementation choices. The Pin1 choreography generated 57 such records, with each carrying structured frontmatter (identifier, title, date, status, the approach it binds, who decided it, what provoked it, what it supersedes and is superseded by, the source files it affects, the evidence available when it was taken). The decision records are distinguished from simple logs by four properties:

##### Scoped

In the Pin1 choreography 43 of the 57 records relate to the entire choreography rather than one approach. The effect of changing any one approach on the other approaches is estimated and catalogued.

##### Attributed

Each record names both the decider and the origin of the decision. The origins can be informative in themselves. For example, 40 decisions arose during implementation, 10 from the PI, 6 from an Adversary audit, and 1 from the original specification.

##### Revisable in part

Each decision is associated with a status: `accepted`, `superseded`, or `partially withdrawn`. The third is needed because one record usually makes several claims, and evidence rarely refutes them all at once. Suppose a record chooses method A over method B because A scored better on some measurement, and that measurement is later shown to be unsound. The choice of A may still stand on other grounds, while the reason recorded for it no longer does. Calling the record superseded would retire a choice that still holds, and leaving it `accepted` would keep advertising a measurement nobody should cite. `partially withdrawn` does neither, and each line of evidence is marked as standing or withdrawn on its own, so a later reader knows which half to trust.

For example, one record in the Pin1 choreography made two claims at once. It reported that the two ways of representing the water around the protein disagree about which candidate molecules stay in the binding pocket, and it reported that two particular candidates had left the pocket altogether. Repeating the work kept the first claim and removed the second. A second run reproduced the disagreement between the two treatments of water, so that finding is now better supported than when it was written. When the two candidates were simulated again under the same conditions, however, both stayed in the pocket: their earlier departure was chance in a single short run, not a property of the molecules. Retiring the whole record would have buried a result that still holds, and leaving it in force would have kept telling readers that two good candidates fall out of the pocket. The record is therefore marked partially withdrawn, with the departure struck out and the disagreement left standing.

##### Rationale

A record says both what turned out to be wrong and why it had looked right when it was written. That second half is what a version-control history cannot supply, and it is what stops the same reasoning being repeated by someone who was not there. The clearest case in this project is the shared control, whose apparent statistical significance was gradually reduced to mere chance over two days once the comparison it rested on was examined; we follow that sequence in §2.8, and the three records that carry it are reproduced in §A.4.

#### Orientation and failure-mode documents

The decision records are written one at a time and never revised, so no single record voices the current state of the project. Two documents are responsible for this and are read at the start of each session. state of the project.md states the goal of the project, what is settled, what has been ruled out and should not be attempted again, what is running, and what to do next. how this project breaks.md lists the mistakes the project has actually made, a count of 21 in the Pin1 choreography. Each entry records the form of the mistake rather than the incident: in almost every case the code took a value out of a table by its position, its name, or a default, instead of checking that the underlying assumptions about the shape of the table were correct. The chosen wrong value was as plausible as the right one, so nothing failed and no test caught it. Fixing one instance does not prevent the same mistake a year later in different code, but recognizing the form of it might, which is why this file is read before any session that writes code. Nine mistakes reached a result and forced a claim to be withdrawn; the rest were caught before they did. Only three of the 21 were caught by an automated check, while the rest were noticed by a person reading output that did not match expectation, which is an argument for writing more such checks rather than for auditing harder (Figure S3C).

#### Process control

Long-running computations (e.g. software builds, optimizations, searches) can be interrupted and cause sessions (and their associated context) to be lost. The ability to recover gracefully from such events without losing previous, valid progress has many obvious advantages. Towards this end, Murmurent is designed to periodically write to file notes that facilitate reproducible recovery including which version of the code, which settings, which input files and which machine were used. When two runs disagree, the system investigates and reports the source of the change (code, settings, data). Before launching jobs, Murmurent estimates the total resources they will require and considers parallelizing the computations. For example, during the Pin1 development Murmurent recognized in advance that a script which docked 16,806 molecules would take several days on a single CPU. It identified a GPU-based variant of the same software, installed it, and distributed the work over the 5 available GPUs, which finished in 7.4 hours of wall clock time. The incident itself was also catalogued in how this project breaks.md as an error to sensitize CC in future computations.

### 2.6 Enforcement of security, privacy, and intellectual-property protection

Security and privacy are concerns in academic settings given the nature of the data, for instance, patient-derived cohorts, unpublished results, and patentable molecules. Traditionally, security and privacy have been maintained primarily by a two-pronged approach: (i) university information technology services, which govern where data is stored and who may read it, and (ii) the judgment of the researchers themselves, who know, for example, not to paste a lead compound into an external service. Such approaches are insufficient to cope with the independence of AAI, which can seamlessly send data to an outside service, without recognizing the disclosure and without prompting the user. AAI is also subject to redirection: studies show that text an agent reads (e.g. a web page, a retrieved document) can steer it toward unintended goals [47, 48] while still appearing to be compliant [49, 50]. Moreover, AAI models themselves are becoming increasingly capable of finding exploitable flaws in software including in the infrastructure that research groups depend on [51].

Murmurent implements protections at four points: beneath the agent, where a guarantee holds regardless of what the agent decides; at the agent, where a dedicated actor reviews what is about to leave; at the boundary between a member and the group, where information passes from a private record into shared memory; and in the identity of whoever is acting. The concrete mechanism behind each safeguard summarized here is given in §A.3.

#### Inescapable rules

Murmurent creates and manages several named directories with specific permission systems in order to minimize the chance that either a user or an agent accidentally deletes precious information. Using so-called *hooks* in CC, which fire automatically at every tool call, the system disallows any attempt to delete or overwrite files placed in the `immutable/` directory. Similarly, only new files may be placed under `append only/`, and existing files may not be deleted or overwritten. Neither depends on the agent’s cooperation, on how the prompt was phrased, or on the user noticing in time. What this reliably prevents is an agent destroying a result by mistake. It is not an operating-system boundary (see §A.3 for more details).

#### Control on what enters shared memory

Murmurent checks the provenance, sensitivity, and authorship of information submitted by users to the Oracle for inclusion in group or institutional shared memory. In the Pin1 choreography, the curated set of published binders and the negative results regarding the pocket’s scoring function represent non-proprietary information that could and perhaps should be promoted to the lab’s Oracle memory, so that others can exploit this information in their studies. Sensitive or proprietary information (e.g. candidate binders identified by the choreography) cannot be promoted.

#### An inspectable log of agent activity

A hook writes one line per agent turn to a log held on the member’s own machine (`agents.log`), recording which agent ran, when, and the short verdict its reply opens with. The member therefore has a chronological record of what each agent concluded without keeping or re-reading the full transcripts, and on a shared lab machine the PI can read the same file.

#### Scheduled reconciliation rather than user vigilance

A read-only security scan searches for secrets, protected health information, and world-readable files, reporting potential violations of security rules. An additional reconciliation routine evaluates users’ files, directories and permissions against the stated lab (or institutional) SOPs, highlighting deviations to the PI. These procedures, which are performed automatically with a regular cadence via the Security Guard, may greatly assist in the maintenance of a secure, well-organized computing environment especially with novice users involved. In the Pin1 choreography, the sensitive information is the candidate structures which represent potential patentable property and must remain local. The Security Guard reviews the output from the choreography including figures, tables, graphic user interfaces, and source code for possible violations.

#### Cryptographic identity, decoupled from access

Membership is asserted through a signed certificate (a *card*): every member, PI, and mayor mints an ed25519-based keypair [52] on first use, and the fingerprint of the public half serves as the unique identifier for the person. A member’s card is signed by their PI’s key and, where a lab participates in a centre, the PI’s card is in turn signed by the centre’s root key, the key at the top of the chain of trust, held by the centre’s administrator (referred to as the *mayor*). A lab that never joins a centre has the PI as its own root (§4.4). Issuance is deliberately decoupled from live authorization: a card attests identity, but access is carried by the registry and by the chat software (Slack here) and GitHub access-control lists, so a valid card grants nothing the registry does not independently confirm, and removing a member revokes card and access together. Taken together, these four protections treat the agent itself as untrusted. Its behaviour must be explicitly verified rather than assumed. We regard this as the appropriate default for any agentic deployment in a research context.

### 2.7 Multi-user operations: groups, projects, and centre administration

Murmurent is organized around four social units: (i) The individual *member* is a single researcher. (ii) A *group* is either a lab, consisting of a principal investigator (PI) and a set of members, or a core facility (e.g. genomics, protein structure, imaging) with a leader and a set of members. (iii) A *project* is a unit of work that brings members together, potentially across groups. (iv) The *centre administration* is an institutional unit, led by the *mayor*, a human role that decides which groups join the centre, holds the centre root key that certifies every PI (§4.4), maintains the directory of groups and projects, owns centre-wide governance and accounting, and supervises cross-unit data transfer. The administrative agents act on the mayor’s instruction and hold no authority of their own (Figure 1B). Onboarding is a bottom-up process: every installer (member, PI, or mayor) runs a one-time setup (`murmurent init`) and declares their role locally. A group need never register with a centre at all and can operate as a standalone, with its PI acting as the root of trust (§4.4).

Projects can form between members of the same lab, or between members of different labs; in the latter case, a project inherits the governance and SOPs of each participating lab. The participating labs keep one official list of a project’s members, and the Registrar issues each member an ed25519-based certificate to prove their membership when challenged. The Millwright agent creates a Git-controlled code repository, a communications channel, and protected file systems (e.g. `immutable/<project>`, `append only/<project>`), and admits only the project members. The Pin1 choreography was run as an inter-group project, with the four approaches developed in parallel by different people and no controller reconciling them.

Finally, the administrative level also manages several agents including a centre-level Security Guard, which audits world-readable paths, secret-shaped strings, and protected storage across the institution, and a centre-level Oracle that maintains institution-wide information. For example, patient cohorts developed by the institution, databases of chemical screens, protected intellectual property generated by labs over time, communal reagents and equipment, and other items with utility beyond any single lab are retained here persistently.

### 2.8 The *Pin1* choreography revisited

We highlight here how Murmurent contributed to the Pin1 choreography of §2.2.

The shared control, described in §2.2, asks each approach to rank the known Pin1 binders above decoys matched to them on size, greasiness and charge. The covalent control first scored an ROC-AUC of 0.815, on a scale where 1.0 is perfect separation of binders from decoys and 0.5 is no better than chance. The adversary objected that decoys matched only on those whole-molecule properties can be told from the binders by their chemical scaffold alone, so the score was measuring scaffold recognition rather than binding. Murmurent made two corrections. It scored each molecule in its bound, post-reaction form rather than its free form, which lowered the score to 0.718. It then required each decoy to carry the same reactive chemistry as the binder it is matched to, which lowered it to 0.537, a score reflecting random chance. Class matching also leaves only two chemical classes in the comparison, against the six required before a result is called, so the final figure records that the ranking was never shown to work rather than measuring how badly it works. All three measurements were reported as underpowered.

A docking score is a weak instrument on a pocket as shallow and solvent-exposed as Pin1’s, and a near-random score still produces a ranked list, so even a top ranked candidate may lead to a false discovery. An independent check on the same machinery makes the point directly. Given Pin1 structures whose true ligand position is already known, the docking program proposes nine candidate positions for each molecule and scores them. The position it scores highest is the correct one no more often than a position picked at random, while the correct position is among the nine it proposed 41.5% of the time, which is within the range published for this type of test. The search finds the answer and the scoring cannot recognize it, which is why the shared control refuses to certify a ranking no matter how the ranking was produced. Murmurent therefore rescored the covalent complexes by molecular mechanics with generalized Born and surface-area solvation (MM/GBSA) [53], a physically independent and more expensive estimate of binding free energy, both from a single minimized structure and across a short molecular-dynamics ensemble. The verdict did not change, and for the same reason: too few chemotypes and too few chemically distinct decoys to support any conclusion. No amount of computation can adjust for the shortage of published Pin1 binders. Murmurent recorded the limit and turned to longer molecular-dynamics runs.

Two distinct obstacles stood in the way. The first is that nothing here was measured at a bench. Every quantity is computed, and sitting in the catalytic site is at best a proxy for inhibition, since a molecule can occupy the site without stopping catalysis. The two covalent approaches *T*_3_ and *T*_4_ are better placed than the reversible *T*_1_ and *T*_2_ for this reason. A molecule that forms a permanent bond to Cys113 blocks the amino acid the enzyme uses to catalyse, so the case that it inhibits rests on chemistry rather than on the docking scores the audit discredited. A molecule that only occupies the site cannot make this argument. Stated as an energy calculation it fails outright, because both compounds known to bind Pin1 covalently are scored badly by the calculation meant to recognize them. Sulfopin, whose bond to Cys113 has been seen crystallographically [37], is outscored by 50 of 80 property-matched decoys, and juglone by 47 of 80; on the corrected interaction energy the two rank 56th and 78th of 80. The covalent route is also narrow: almost all the clean and selective precedents use one warhead chemistry, meaning one reactive group, so the range of reactivity Murmurent accepts is a rough filter for triage rather than a hard criterion.

The second obstacle lies in the measurement rather than in the design of the choreography. The free-energy rescoring treated water as a featureless continuum rather than as individual molecules, which is the cheaper approximation and the less faithful one. Murmurent repeated the dynamics in explicit water, and the two treatments disagreed almost entirely about which molecules stayed in the pocket, which retired a set of claims that had rested on the cheaper model. Explicit water did not rescue the ranking. Repeating the free-energy calculation itself in explicit water is the one clear increase in physical realism that these results do not rule out.

Murmurent built and tested the four approaches iteratively against the same criteria, and held long-running, interdependent calculations together over several days. All four produced a shortlist, and the shared control certified none of them. That is not surprising, since docking approaches are weak on such shallow pockets [32], and no compound reported here has been synthesized or assayed. Nevertheless we had at least two significant successes. First, our medicinal chemists picked out several promising candidates from the 25 best hits across the approaches, and said the exercise had shown them where each of the four is strong and weak; that led to changes which improved the choreography and surfaced further candidates. Second, the refusal to certify is important. Without a shared control and a common evaluation across the four approaches, a ranked list of top candidates is easy to trust, and wet-lab effort would have been wasted chasing false positives.

## 3 Discussion

Murmurent’s most immediate contribution is to lower the threshold at which a biomedical researcher can bring computation and data analysis into day-to-day work. A new member inherits a working set of agents rather than building their own, and inherits with them the memory and traceability arrangements that make a session start from established knowledge of the lab. A wet-lab trainee who would previously have needed a sustained collaboration with a biostatistician/bioinformatician can now carry at least some analyses themselves through ordinary conversation. This does not reduce the importance of quantitative scientists involved in a project but may reduce the load placed on them. A PI with little information-technology support can easily describe how they want data, code and analyses organized, and Murmurent will uphold this vision with hooks and agentic review. This approach is far more efficient than repeatedly reminding people, especially those early in training, who are the ones most likely to make an irreversible mistake.

Multi-user support and the tiered memory system provide a vault of knowledge that is easily shared across group members and across projects. This knowledge includes

- qualitative knowledge (build design decisions, observations, near-misses, and the reasoning behind a methodological choice) to assist CC when it modifies an existing build or adds a new approach to an existing choreography;
- intermediate computational data objects (compound screens, parameter sweeps, cohort-level summaries, instrument-specific calibration histories) which are typically highly project-specific information that could help direct how additions and extensions to a project are carried out;
- sensitive data (patient cohorts developed with partner hospitals, locally curated assay libraries), which cannot be deposited publicly but must remain queryable across years and personnel changes.

The Oracle’s contribution is to record these three types of knowledge with explicit provenance, sensitivity, and audit, alongside the data-governance invariants that protect them.

Academic centres may also carry some constraints that are not always present in industrial settings. Dedicated information-technology and bioinformatics resources are typically limited; Murmurent’s agents and toolkit allow groups to share the cost of developing that infrastructure. Academic centres also carry a teaching mandate. The review agents, inspectable audit logs and security/privacy enforcement together provide a robust set of teaching tools to introduce non-bioinformatics personnel to the power of AAI in data science. EDID and SGBA+ review are not peripheral to academic biomedical research within Murmurent: the Conscience agent is an active structural component of day-to-day research increasing mindfulness and better ensuring respectful, inclusive communications. In the context of lab meetings and research-discussion groups, we find that CC’s capacity to create GUIs and present these results “live” makes it a powerful translational tool, greatly facilitating understanding and debate.

Murmurent is not in competition with lab-in-the-loop platforms, self-driving laboratories, and multi-agent scientific-reasoning systems. Each addresses a specific scientific end-point with a vertically integrated infrastructure, but Murmurent addresses a different problem: the coordination, governance, and memory of academic research across multiple end-points and disciplines. Robin [20], CLADD [12], or an AI co-scientist [21] could easily run within a Murmurent choreography. The same observation applies to the rapidly growing collection of techniques for specializing AAI in biotherapeutic discovery (e.g. domain fine-tuning of chemistry language models, curated retrieval indices over assay and bioactivity databases, structure- and graph-aware molecular encoders, and tool-using agents specialized for retrosynthesis, docking, or property prediction). The substrate Murmurent provides is intended to absorb these as MCP wrappers, so that the group benefits from advances in agentic methods research without each lab having to integrate them in isolation. The reference implementation is built on CC (Anthropic), but each component of Murmurent should be implementable in other AAI systems.

Although Murmurent likely lowers the bar for new bioscience researchers to exploit AAI, it remains undetermined whether this is sufficient for individuals who may have little or no programming experience or statistical background. There are many concepts, perspectives, and approaches that quantitative scientists learn that subtly affect how these individuals interact with an AAI. For example, quantitative experts have an understanding of what is or is not computable, how complex a software build may be, what efficiency means in computational settings, and may have intuitive insight into the mechanisms of a particular statistical test. Users of Murmurent without such a quantitative background may still have considerable difficulty finding the correct level at which to build prompts, using phrasing that the underlying AAI will understand, and may lack an appreciation of modular construction and piecewise improvement.

Data security and privacy are not guaranteed by our current safeguards. The rules and hooks implemented in Murmurent protect against agents directly uploading sensitive data to an external website, but they do not stop a script written by an agent from doing so. Moreover, we cannot rule out the possibility that the search techniques used by the Security Guard fail to identify sensitive data in some instances, or that data sent to a third party contains subtle information that could be exploited. We view Murmurent as a first step in the process of using AAI to assist with security and privacy.

## 4 Methods

All of Murmurent is open source but requires CC, which provides a command-line interface to Anthropic’s large language models. An installation delivers five things: (i) the reference agents; (ii) the hard rules and the choreography patterns; (iii) the Murmurent command-line tool; (iv) the guard programs and the service adapters through which agents reach outside tools and institutional systems; and (v) a local dashboard. Reference agents, hard rules and choreography specifications are all markdown files. The guard programs, the service adapters and the Murmurent command-line tool are written in Python 3.12 [54], with small utilities in Bash. Personal and lab knowledge is held in Obsidian-compatible markdown vaults. Context passed between agents and tools uses MCP. Murmurent running within the VSCode editor provides a four-pane launcher built on tmux, a terminal multiplexer that keeps several sessions side by side (`scripts/open murmurent.sh`).

The dashboard is the visual counterpart to the command line interface (CLI), and gives three different views. At the level of the user’s machine, it displays their repositories, project membership, their personal Oracle and their daily lab notebook. At the level of the group it shows the members, the projects, and the Lab Oracle. For the mayor, it shows the centre, with its participating labs and cores and any join requests awaiting a decision. All actions offered by the dashboard have equivalent CLI versions.

### 4.1 Skills and choreographies

Murmurent distinguishes two levels of automation. A *skill* is a single command a user can invoke that carries out one self-contained action; for example /murmurent-push saves and uploads the current state of a Murmurentaware code repository. Skills are an established construct in CC. A *choreography* is a written, multi-party pattern combining skills, agent invocations and human approvals into a coherent whole. It is the level at which group and centre work is organized, and it is how Murmurent coordinates work without anyone directing it centrally.

### 4.2 Agents

The default Murmurent defines 13 reference agents (§2.3), each an ordinary markdown text file (`agents/*.md`) installed into the local CC configuration when Murmurent is set up. A table naming each agent, the social unit it acts for, and the words its verdict is drawn from is kept with the software rather than here, so that it stays correct as agents are added. A member may create *ad hoc* agents for their personal work (through the dashboard, the CLI, or by writing their own markdown file) and keep them in their own vault and GitHub account. The agents stay in that member’s environment and are not shared. When a centre is defined, the administrative level acts above any single group via several dedicated agents: the Registrar, which keeps the centre roster and issues membership certificates, and the Centre Millwright, which compares the state of shared infrastructure (file permissions, Slack and GitHub membership) against that roster and corrects any differences. That comparison (`murmurent reconcile`) can be scheduled to run daily. It reports what it finds and changes nothing unless it is asked to repair, so a difference between the roster and the infrastructure is raised with a person before anything is altered.

Three agents are dedicated to verifying user identity (§4.4). The Registrar issues the cards that certify PIs and publishes the centre’s public signing key and its list of withdrawn cards. The Millwright and Security Guard together keep keys in good order, checking card status against live Slack and GitHub membership and checking that private key files are not readable by anyone else.

### 4.3 Hard rules and data governance

Four rules govern how an agent may act, subject to the limits set out in §A.3. They govern work done through Murmurent rather than an arbitrary process touching the same filesystem. The rules are: (i) nothing under `immutable/` may be altered by an agent; (ii) files under `append only/` may be added but never replaced, with successive versions numbered; (iii) every Oracle entry carries a checked header recording title, date, project, sensitivity, tags and sources; (iv) an agent’s reply must open with its verdict. The four are not upheld in the same manner. Rules (i) and (ii) are enforced by two small programs, `raw guard` and `protected paths`, which run before any operation an agent attempts that could write to or delete a file and refuse those that would break either rule; nothing an agent decides can evade them. Rule (iii) is checked when an entry is submitted, so a malformed header is refused at that point rather than when it is read. Rule (iv) is an instruction carried in each agent’s own definition and is not enforced by a program at all.

### 4.4 Identity

Membership in Murmurent is carried by a signed certificate, called a *card*. Identity is established from the bottom up. After installation each user runs one short setup command (murmurent init) that records whether they are a member, a PI or a mayor, so a lab can adopt Murmurent and work on its own long before any centre exists to register with. Every member, PI and mayor creates a signing key pair on their own machine the first time they run Murmurent (murmurent identity-init), using the ed25519 signature scheme [52]. The Millwright sets up everything else a new member needs: their SSH keys, a working copy of the repository, the layout of their Obsidian vault [43], and the paths their tools will use. The private half is kept in a file readable only by its owner, under ∼`/.murmurent/keys/`, and is never copied or transmitted; a short fingerprint of the public half is the holder’s unique identifier. Before a PI or mayor writes a prospective member’s fingerprint into a card, that member has to prove they truly hold the matching private key. The issuer sends them a short one-off phrase, and they sign it and send it back.

The chain of trust follows the organization it certifies. At the top of any such chain sits a *root*: a signing key that nothing above it vouches for. By default a lab stands alone and its PI is that root, issuing their own card (`murmurent pi-init`) and signing member cards directly. If and when a lab later joins a centre, the mayor’s centre root key also signs the PI’s card. The member cards themselves do not change, yet they now verify up to the centre as well: the chain gains one further link above the PI. The software enforces the shape of the chain: only a root may sign a PI’s card, only a PI may sign a member’s card, and a member’s card cannot sign anything at all. Every signature is made the same way, using one method and one format, which is fixed in advance rather than agreed when a card is checked (§A.3).

A centre makes four things public in the directory of installations (maintained at the GitHub repository murmurent public): the key it signs with, the key others use to encrypt messages to it (age, https://age-encryption.org), its current list of cancelled cards, and its own entry in that directory, which it signs itself. Each member records which key the centre signs with the first time they see it, having checked it through some other channel such as a phone call. If the published key later changes, the check fails rather than quietly accepting the new one, and somebody has to find out why.

Signing PIs’ cards and signing the list of cancelled cards are the only two things the centre’s root key is ever used for. No automated process holds it, every signature it makes is one the mayor performs deliberately, and its single backup is encrypted and kept off the network. § A.3 explains why this key is treated differently from every other one.

The same mechanism applies to individual projects. A project card binds a member’s existing key to one project rather than to the lab as a whole: it names the project alongside the lab, and gives the holder the role of project member. In every other respect it is an ordinary member card chaining through the PI to the root, so existing checks accept it unchanged. Project cards have their own list of cancellations, kept separately from the lab’s, so deleting a project cancels every card issued under it in one step and leaves everyone’s standing in the lab untouched. One file per project records which members hold a valid card for it, together with where the project’s code is kept and which chat channel and GitHub repository belong to it; one file per member records that member’s card fingerprint, card number, email address and GitHub login. These files are the authority, and any other record of issued cards can be rebuilt from them.

### 4.5 Oracle memory and MCP retrieval

The personal Oracle is the `oracle/` folder of each researcher’s vault, a plain folder of markdown files readable in Obsidian (Tier 2; §2.4), holding one file per entry, each with a checked header. A small adapter (`oracle server.py`) lets agents search the Oracle and lab-notebook tiers by project, date, tag, sensitivity and source. Entries that have been reviewed are promoted to the Lab Oracle, which agents may read but not write, through the `murmurent oracle publish` command, and that command refuses any entry marked `restricted` or `protected`. The same vault holds a daily lab-notebook tier of dated entries that need not carry the Oracle header, which the search adapter looks through alongside the personal and lab tiers on request. It also holds a `maps-legends` area that each member arranges to suit their own navigation and that Murmurent does not check, although reference agents read it before writing so that tags and structure stay consistent as entries accumulate. The vault’s fourth kind of content, `murmurent data/`, holds reference material of any sort, including PDFs, spreadsheets, protocols and images, which agents read as needed and which is not checked against any schema. There are two ways to read it: (i) an agent that already knows which file it needs reads that file, or (ii) an agent that does not know what is there asks a small service to list the contents of the folder, and then asks it for whichever item it wants (`data list` and `data read`).

## 5 Declarations

### 5.1 Ethics approval and consent to participate

Not applicable.

### 5.2 Consent for publication

Not applicable.

### 5.3 Availability of data and material

The reference implementation of Murmurent is available at https://github.com/hallettmiket/murmurent, or can be installed from the Python Package Index as murmurent (https://pypi.org/project/murmurent; version 2026.9.8 at the time of writing). Developers can access the development version at https://github.com/hallettmiket/murmurent_dev. The public onboarding hub, which holds the institution directory and a list of public choreographies is at https://github.com/hallettmiket/murmurent_public. The Pin1 choreography can also be cloned directly from https://github.com/tt8804/inhibition_public, tag v1.0-paper. All Murmurent components are released under the Apache license version 2.0.

### 5.4 Competing interests

The authors declare no conflicts or competing interests for any aspect of this manuscript.

### 5.5 Funding

We acknowledge financial support from the Natural Sciences and Engineering Research Council of Canada to MTH and VD, and the Canada Foundation for Innovation (CFI, #43481) for the computing infrastructure (GG, VD, MTH). MM and BK received the Ontario Graduate Scholarship, provided by the Province of Ontario and Western University. BK and YX received the TBCRU MSc fellowship, provided by Breast Cancer Canada. TSB and DRE received funding from Genome Canada and Ontario Genomics via the Genomic Applications Partnership Program (GAPP). KPL and XZZ are supported by the Canadian Institutes of Health Research, Ontario Institute for Cancer Research and Canada Foundation for Innovation.

### 5.6 Authors’ contributions

VD, MTH conceptualized the project. TW, TSB, MM, NHS, EJM, YX, BK, VG and MTH developed the code underlying Murmurent. TW, HEE, TSB and MTH developed the code underlying the Pin1 choreography. AJ, AM, LW and NHS designed, built and experimented with the Conscience agent. TW, HEE, NHS, MTH wrote the manuscript. MM, EJM, LW, and VD provided critical feedback. SLB, DRE, GG, KPL, XZZ, VD and MTH provided financial support.

## Appendix A

### A.1 The Teacher agent

The Teacher returns one of exactly two verdicts, and each is a definite claim rather than a grade. *Explained* asserts that the mechanism has been delivered in plain terms and that none of the five conditions set out below was triggered. This asserts that the agent can confidently support its claims. *Gap* indicates a failure but with some information: the agent declines to explain the target topic, names which of those conditions it failed, and quotes the particular step the reader would otherwise be taking on trust, together with what should be read instead. There is deliberately no third verdict between them.

A *Gap* is triggered by any one of five conditions:

- nothing was read: a debrief with no transcript, or an account of a paper that was never opened;
- the causal chain breaks, in that the agent, rebuilding the mechanism from memory rather than reading it back from the source, reaches a step it can only assert;
- the counterfactual cannot be completed: unable to finish *this would have come out differently if `X`*, the agent has described a sequence of events rather than a reason;
- the jargon budget is exceeded, that budget being three technical terms, each defined where it first appears;
- no quiz question can be written whose wrong answer names a specific misunderstanding, which applies to rendered pages only.

The last of the five is the sharpest, because a plausible wrong answer cannot be written by someone who does not already hold the right model. The output conventions are as follows:

- a reply opens with its conclusion in plain bullets, so that a reader who stops there still has the point;
- detail is scaled inversely to the reader’s expertise, on the view that over-explaining to a competent colleague is a harm rather than a courtesy;
- at most one load-bearing analogy is used, and the point at which it breaks is stated;
- compression is applied to the language but never to the uncertainty, so that a caveat carrying real doubt survives it.

The agent is deliberately narrow, and its boundaries are drawn at the other shared agents (§2.3). For example, it does not render visuals such as figures or diagrams, but instead hands such tasks off to the Artist. The need for persistent memory is handled via the Oracle. Literature and database sources are deferred to the Bookworm, which means that readings will remain inside the centre’s data-governance boundary rather than on the open web.

As a technical side note, the Teacher’s Course mode is not carried out by the Teacher agent. When the Teacher recognizes that a learning task may require a sustained dialogue with a student, the Teacher decides to return *Gap* and names the course skill, which the calling session then loads. This is because a course must interview the learner and carry state between sessions, which is beyond the capabilities of the current subagent.

Debrief mode reads the transcript of the session that invoked it. This is the one capability that gives the Teacher sight of material the rest of the agents do not see, and it is fenced accordingly (§2.6). The agent may read only its own invoking session, resolving the path by derivation and refusing any request that names a different one; it never enumerates the directory holding transcripts, which on a shared machine carries every session run on it; it quotes reasoning but never tool results or attachments, so that what a step did is described without reproducing what it saw; and in a project marked restricted it quotes nothing at all.

**Fig. S1.**
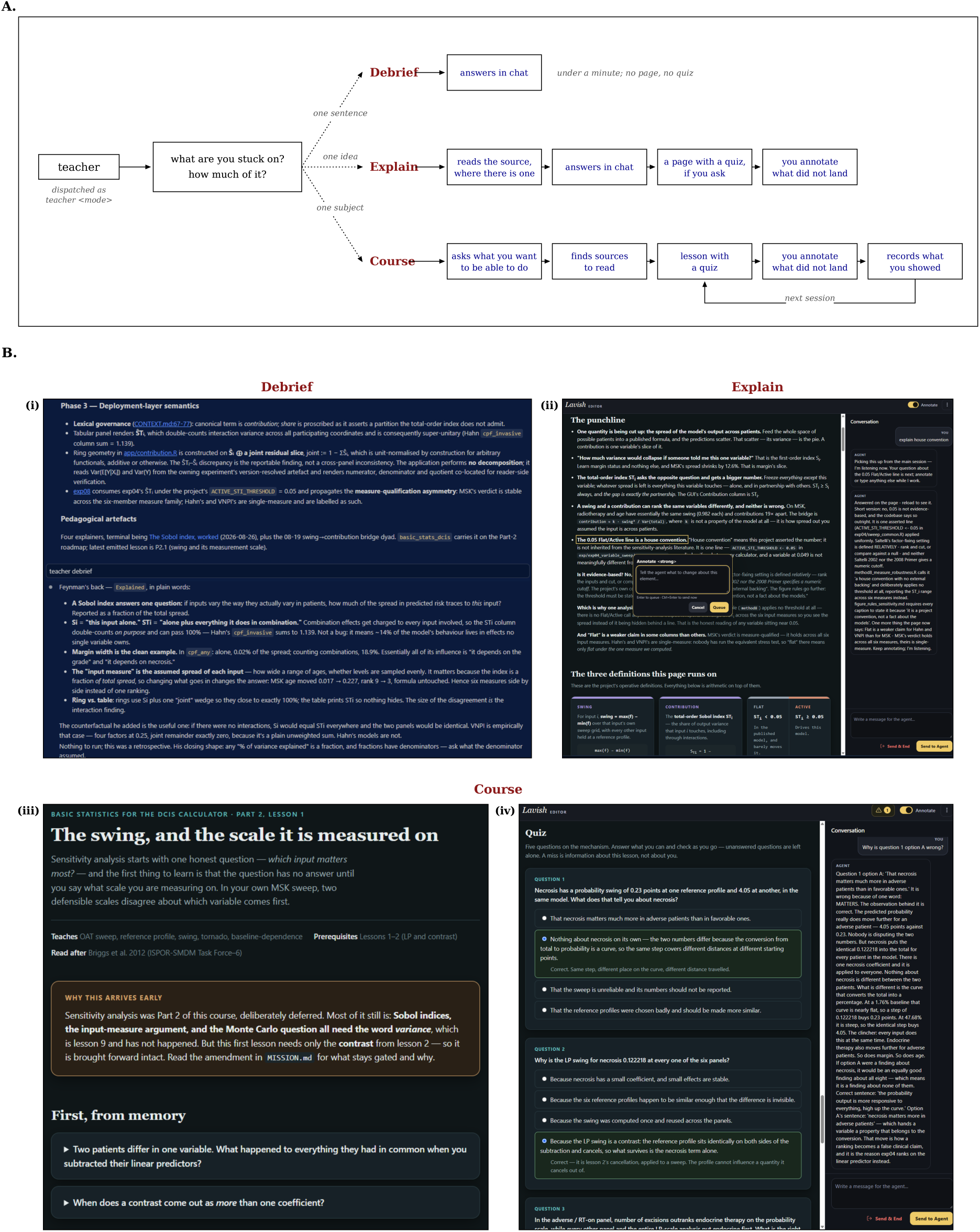
The Teacher agent and its three modes. **A.** The scope of what the user wants to understand determines the mode. **B.** (i) *Debrief.* Re-phrases a single CC sentence in plain words restricted to a budget of only 60 seconds. Output is in the chat window of CC. (ii) *Explain.* Provides an explanation for a chosen concept or source, producing one page of output in addition to a self-grading quiz on request. The user can simply highlight text in CC. (iii) *Course.* Via the underlying murmurent-course skill, the Teacher builds a multi-component course to systematically explain a chosen topic to the user with (iv) an accompanying quiz.

### A.2 The Conscience Agent

**Fig. S2.**
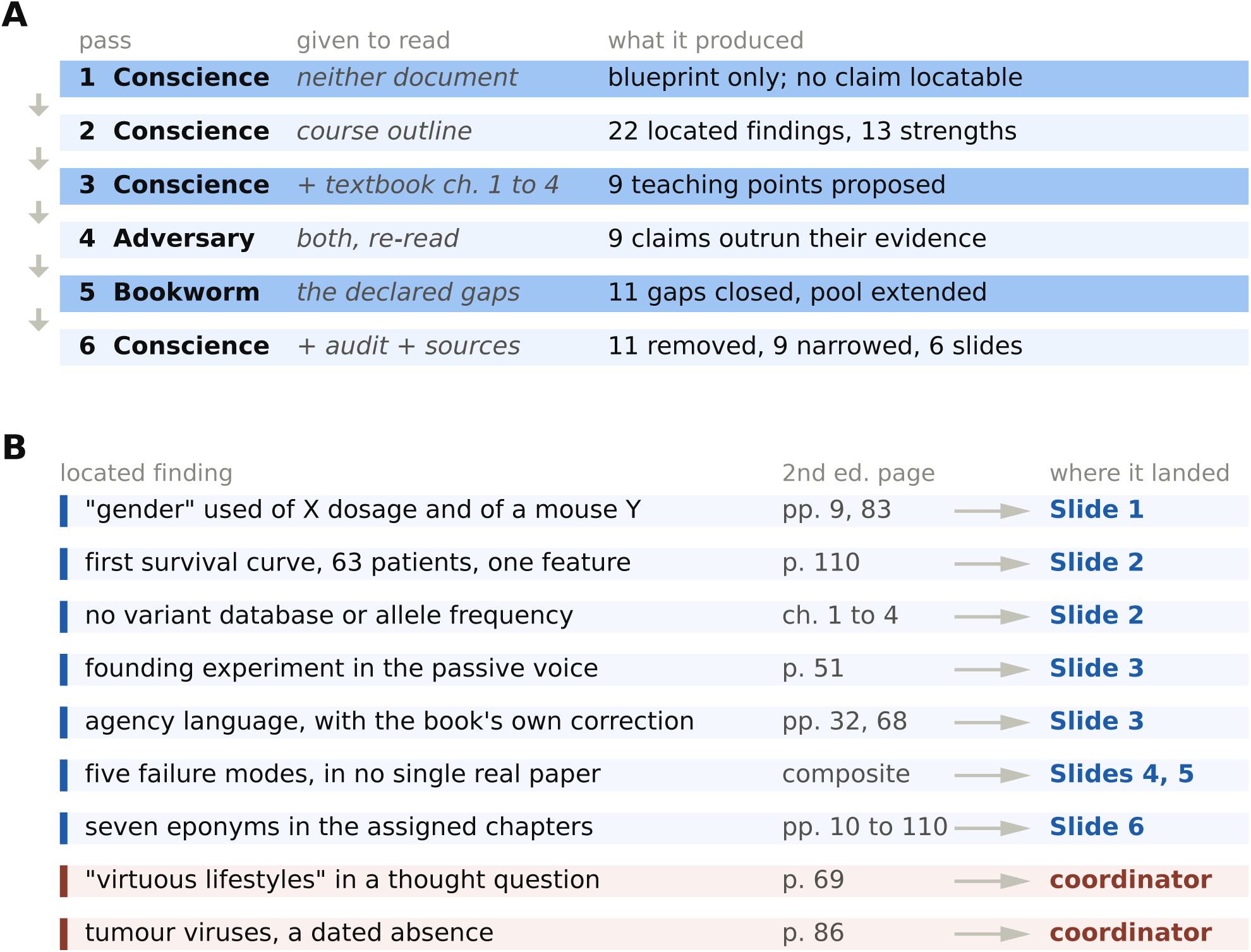
Studying the Conscience on a teaching task. **A.** The sequence of agents over one course, the documents each pass could read, and what it produced. The Conscience opens, the Adversary audits its output against the same sources, the Bookworm closes the gaps the Conscience had declared but could not fill, and the Conscience then rewrites against both. **B.** How the findings chose the slides. Each finding is located on a printed page of chapters 1 to 4 of the assigned textbook. A finding that falls inside the five lectures this instructor teaches becomes slide material; two findings that fall outside them are real but unplaceable here, and are routed to the course coordinator instead.

We analysed the behaviour of the Conscience agent on a task related to the development of teaching material. Specifically, we asked the system to assist with the preparation of the first five lectures in a senior undergraduate biochemistry course which covers the fundamentals of molecular oncology via the standard textbook *The Biology of Cancer* by R. Weinberg [55]. The prompt had three parts. First, identify places in the syllabus [56] which do not follow best practices with respect to EDID and SGBA+. Second, read chapters 1–4 of the textbook and find the places where an instructor could provide students additional EDID and SGBA+ information to complement these topics with special emphasis on the Ontarian and Canadian context. Third, it was asked to create approximately five slides to cover these issues and also provide one worked example of a molecular oncology finding in which SGBA+ was not considered, with the explicit instruction that the example be anonymized (to avoid criticizing any specific author).

#### How the agents ran

After reading both documents, the Conscience identified 22 items in the outline and 9 key places from the textbook (Figure S2A). The Adversary then re-read the same two documents and audited every quotation, count and inference in the Conscience’s output, identifying several issues (described below). The Bookworm searched the literature for each such issue, and added its findings (and non-findings) to the shared resource pool. This included reading several manuscripts from the primary literature referenced in the Weinberg textbook which provided a capacity to better evaluate the Conscience findings. The Conscience then withdrew what the audit refuted, narrowed what it had overstated, and wrote the slides from what was left.

#### The course outline

The system identified numerous (22) issues with the syllabus including several related specifically to EDID and SGBA+ which had relatively easy fixes:

- provide a means for students to declare what name and pronouns they use, and
- provide a content note that the course covers topics of cancer mortality and inherited risk, which could be significant issues for some students.

Other issues raised are more systematic in nature, requiring careful consideration and structural changes. For instance, other than the course outline acknowledging Indigenous peoples in the preamble (specifically the land acknowledgment and mention of the National Day for Truth and Reconciliation), the proposed curriculum and method of student evaluation did not incorporate any aspects related to EDID and SGBA+. For example, none of the four learning outcomes asks which populations a body of evidence came from, and across 14 weeks of mechanistic content there is no point at which cancer outcomes are allowed to differ between populations, so the biology is presented as ‘universal’. The Truth and Reconciliation Commission issued Calls to Action on both curriculum and health that speak to exactly this. Ontario has published material that would answer them. One example is a report on cancer in First Nations people co-produced by Chiefs of Ontario, Ontario Health’s Indigenous Cancer Care Unit and ICES, under a data-sharing agreement implementing OCAP®, the First Nations principles of ownership, control, access and possession. There was the additional suggestion that the assigned paper project (worth 45%) would be a good mechanism to allow students to address issues related to Indigenous peoples.

#### The textbook

Several EDID/SGBA+ issues in the Weinberg textbook were highlighted by our system (Figure S2B). The word *gender* appears only twice in 130 printed pages, and both times it is used where sex is intended. Both passages are about chromosomes, and neither is about the social roles the word denotes. In the first case it describes X chromosome dosage, which is a count of how many copies of a chromosome a cell carries, and in the second it describes the Y chromosome of an inbred mouse strain, although there is no evidence that mice have a concept of gender. An end-of-chapter thought question asks what fraction of cancers is avoidable *through virtuous lifestyles*, placing a moral term inside a question about causation. Weinberg almost certainly intends ‘virtuous lifestyle’ wryly, as shorthand for the familiar list of lifestyle factors, and expects the student to unpack it. The founding experiment (tumors are clonal) of chapter 2 is introduced in the passive voice, with neither the reporter nor the provenance of the tissue stated.

The Conscience agent also acknowledged that the Weinberg textbook did many things well. For example, in one place the book calls cancer cells “renegades with an agenda”. This indicates that the Conscience agent is wary of language to the effect of the “war on cancer” and other highly poetic analogies which Weinberg has successfully avoided using throughout all chapters. The textbook in fact clarifies this position just two paragraphs later.

#### Discussion point slides

The system decided on six slides that follow the issues identified in Figure S2B. The slides themselves are reproduced in §A.5. Each slide is listed below with annotations made by three specialists in EDID in the biosciences.

<u>Slide 1</u> examines two common bad writing habits: treating sex and gender as one variable and reporting race as biology with no named mechanism. *Specialist commentary (co-authors LW, AJ, AM):* Treating race as biology is the surviving edge of scientific racism, a pseudoscience that classified people as superior and inferior, concepts that underwrote eugenics. The slide also states that neither sex nor gender has only two values and that neither is fixed for life, which is right and abstract. The values should be named, and the list should include ‘two-spirit’.

<u>Slide 2</u> makes one point with two examples: a claim built from one group of people gets applied to everyone. The first example is the 63-patient survival curve from page 110. The second is the reference databases used to judge whether a genetic variant is dangerous, which are drawn mostly from people of European ancestry, so a variant that is ordinary in an under-represented population appears as rare, and rarity is taken as evidence of harm. Unlike the other findings, these examples are errors of omission: the textbook does not say something incorrect but simply does not raise the issue. *Specialist commentary:* This is an important topic as it touches upon the Euro-Western convention of treating the white, able-bodied male as the default research subject.

<u>Slide 3</u> pairs the passive voice with deficit framing and is built on the chapter 2 sentence above and on the book’s own agency language, quoted with the hedge and the correction the book supplies. *Specialist commentary:* This ties to issues of informed consent, and the asymmetry of class and race between whoever took uterine tissue in 1965 and the Black women it was taken from.

<u>Slides 4 and 5</u> provide the single worked example of SGBA+ issues going unaddressed in molecular oncology. To ensure anonymity, this is a composite example.

<u>Slide 6</u> discusses the use of eponyms in cancer (and other diseases). This was chosen because the target chapters had several examples of eponyms including one example that originated from Western University. *Specialist commentary:* The lymphoma is named for an Irish surgeon serving in the British colonial medical service in Uganda, which makes the eponym itself a record of colonial medicine.

### A.3 Security measures

We expand on the mechanisms behind each safeguard discussed in §2.6, and on what each does not do.

#### Two rules about files, and the limits of what enforces them

Before any command an agent issues reaches the disk, Murmurent inspects it and refuses it if it would change a file under `immutable/`. A second check applies the matching rule to `append only/`: new files may be added, but files already there may not be overwritten or deleted. The check reads only the command itself, so a command that starts a script is judged on that one line, and any write performed by the script itself goes unseen. The check also belongs to one person’s own installation and runs only while they are working through an agent, so it does nothing in an ordinary terminal, nothing for a program run outside a session, and nothing for a colleague who has not installed it. It prevents an agent destroying a result by mistake.

#### Why file permissions, and not the agent, decide who may write

Since the agent safeguards do not provide operating system security, the same rule is applied a second time by the file system itself, on the principle that an agent should not be the only thing guarding its own access. Each project is given its own user group, and that group is allowed to read `immutable/` and `append only/` but not to write to them, so a program that never goes near an agent still cannot overwrite a result. The Security Guard agent then compares those permissions against the centre’s own list of members, so whichever agent grants access is not the only one checking it.

Setting file permissions requires administrative rights on the machine that holds the data, and a centre frequently does not have them. On a national computing facility, on storage run by a university’s IT department, or in cloud storage, permissions are somebody else’s to set. Murmurent then only reads them: it compares the permissions it finds against the centre’s list of members and reports any difference to whoever does administer the machine. The second pair of eyes survives even where the power to enforce does not.

#### Identity: the purpose of the card design

Every card in a centre traces back to a single signing key held by the mayor, which signs each PI’s card, and each investigator in turn signs their own members’ cards. A card is checked by following those signatures back to that common origin. Requiring a member to sign a one-off phrase before their key is written into a card stops an issuer vouching for a key the requester does not control. Fixing the signing method in advance, rather than agreeing it when a card is checked, denies an attacker the move of offering a weaker method and having it accepted. Requiring the shape of the chain means a stolen member key cannot issue further cards, because a member’s card may not sign anything.

The signing key is called a *root* because nothing above it vouches for it, and a machine trusts it only because its fingerprint was confirmed through an independent channel the first time that centre was encountered. It fails in two quite different ways. Losing it halts issuing and cancellation until a replacement is distributed, which is an availability problem and recoverable. Leaking it lets an attacker impersonate the centre, which is not recoverable, and that asymmetry is what justifies keeping the single backup encrypted, off the network, and away from any automated process.

Withdrawing access and cancelling a card happen in a fixed order: access is withdrawn first, because that is what actually stops someone working, whereas a cancellation takes effect only as other machines fetch the updated list. Cancelling first would leave a window in which a member had been formally removed and could still work. A card whose cancellation list cannot be fetched is refused rather than accepted, since otherwise blocking access to the list would silently reinstate every cancelled card.

#### Restricted material cannot reach shared memory by the ordinary path

Every Oracle entry carries a sensitivity of `standard`, `restricted` or `protected`. `standard` is the ordinary tier and may be shared with the group. `restricted` covers material that is not yet the group’s to hold, such as an unpublished result or a candidate molecule with a patent position to protect. `protected` covers material a data-use agreement or research ethics board keeps inside the member’s own account, patient-derived records being the usual case. The command that promotes a personal draft into group memory refuses any entry marked `restricted` or `protected`, so such findings stay in the contributing member’s own vault. The tiers name how an entry must be handled rather than what it is about, which is deliberate: a label that names the subject invites a reader to guess the handling from the subject, and the two do not always agree.

#### Onboarding without a public record

A prospective member encrypts their join form to the centre’s public key and emails the ciphertext to the published join address, where the mayor decrypts it locally. The public directory of centres holds only a contact address and each centre’s public key, and a centre appears in it only when its mayor deliberately publishes its row. No member-identifying information is stored publicly under this flow: netnames, hostnames and data paths are exchanged only after the registrar engages with the requester directly.

#### What this design does not provide

Access to the registrar dashboard rests on a secret shared among the operators trusted to run it, which establishes that a caller belongs to that group but not which member of it they are. Each change to the registry is recorded against the handle an operator supplies, and because that handle is asserted rather than verified, the audit log attributes actions only as reliably as operators identify themselves. Per-operator credentials, or authentication through an institutional identity provider, would remove this limitation and are left as future work.

### A.4 Decision records, the failure catalogue, and run manifests

§2.5 states that three kinds of document outlive both the session and the person. We found that these documents were instrumental in the design of choreographies. This section shows more concretely the role of these documents in the Pin1 choreography.

#### Decision records

A decision record is created for each consequential choice. A record is warranted if and only if a choice would be expensive or confusing to reverse, or when a fact in the code or analysis would be hard to decipher by a user several months later. Routine implementation choices do not warrant a record. Each begins with a header in a fixed format, followed by prose in three parts: the context that forced the choice, the choice itself, and its consequences. The header facilitates search queries.

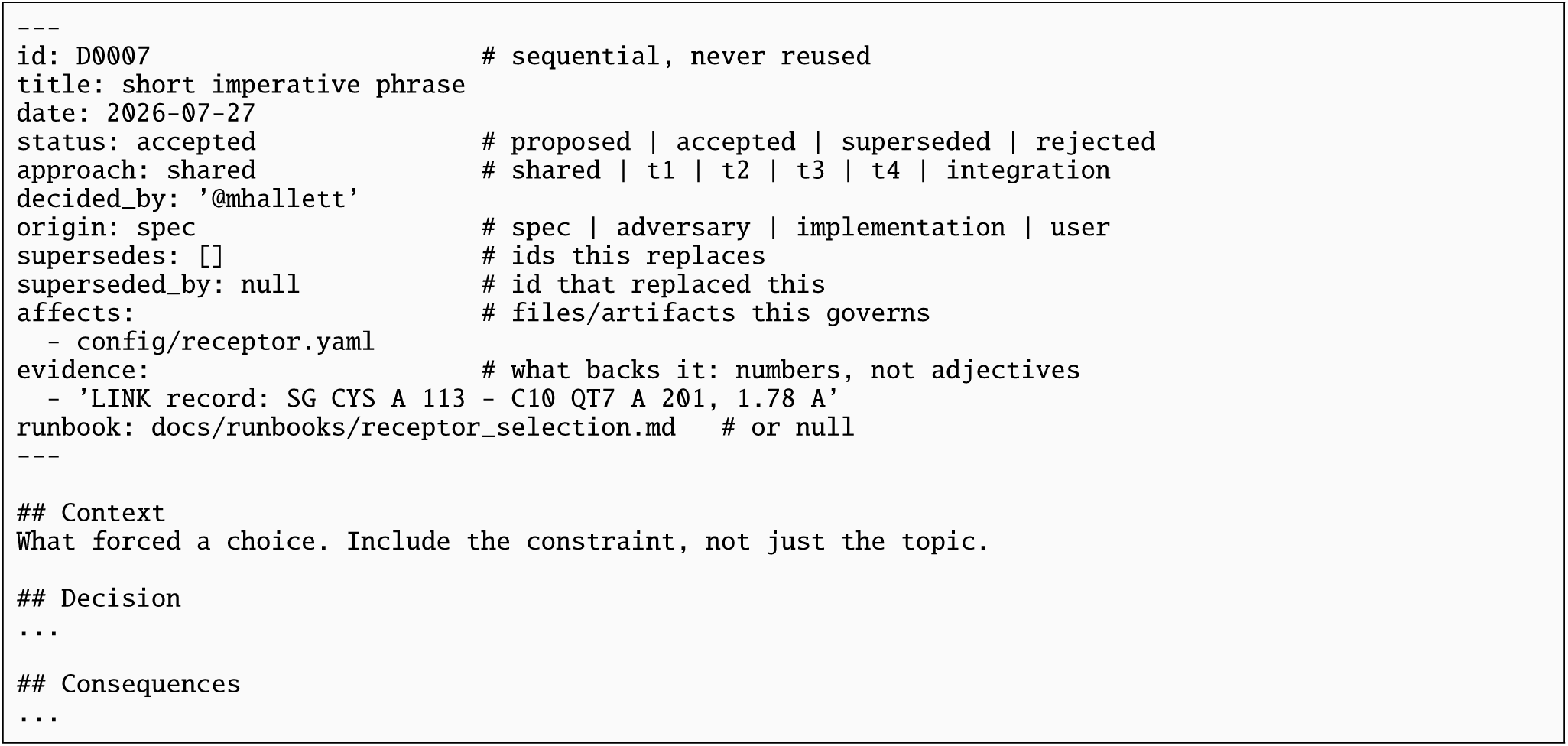

The three records below are the audit sequence described in §2.8, in which the apparent statistical significance of the covalent control is reduced to mere chance over two days. Together they show a claim being withdrawn openly by the system, with the evidence for each step attached to the step that used it.

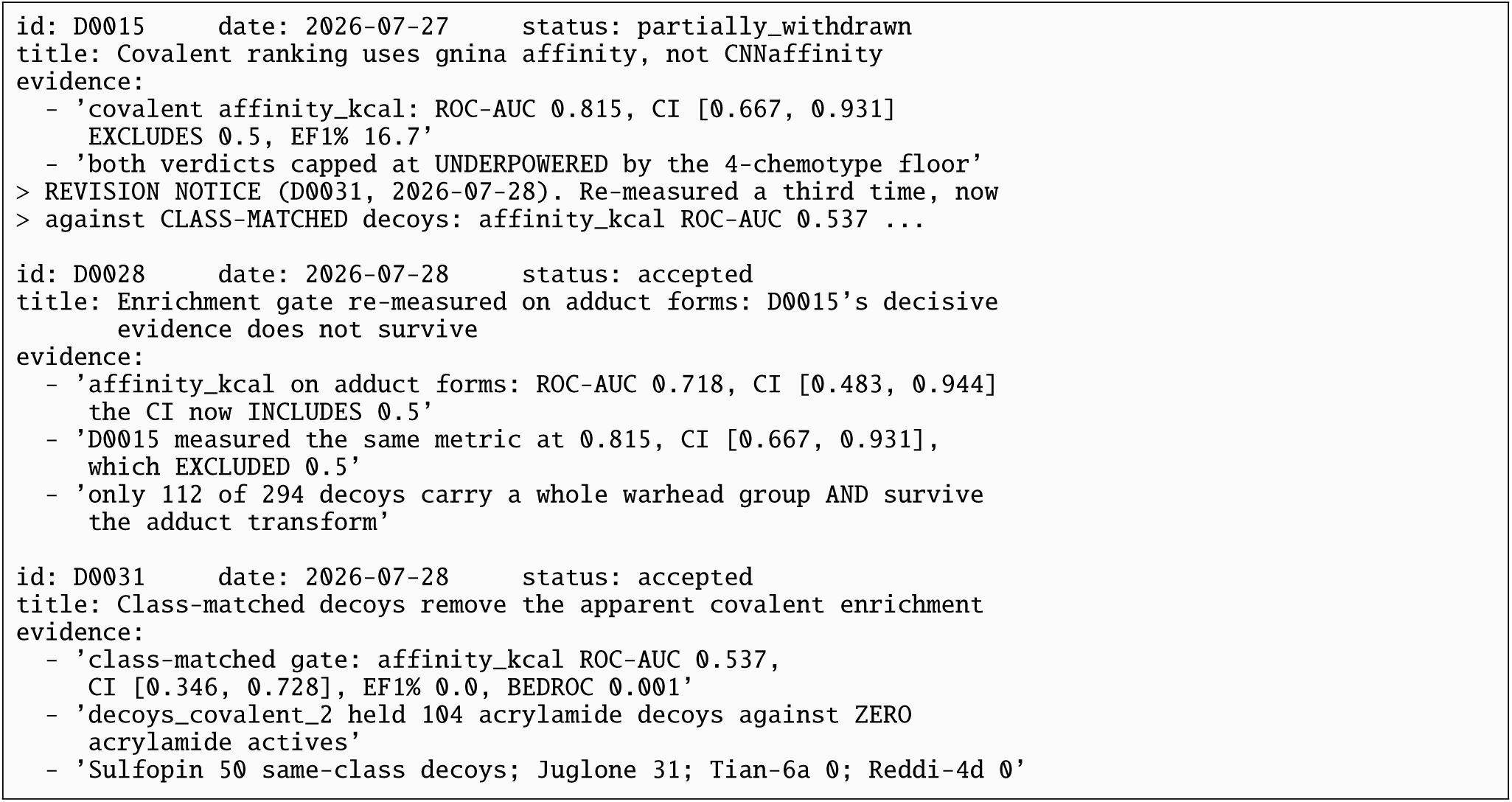

In the evidence fields above, EF1% is the enrichment factor in the top one per cent of a ranked list, meaning how many times more known binders appear there than chance would put there, and BEDROC is a related measure that weights the very top of the list most heavily. Three properties of the decision set are worthy of further note (Figure S3A). First, most records bind the whole choreography rather than one approach, which is what keeps four independently built approaches comparable at the end. Second, every header carries an evidence field listing what supports the decision, and the rules require those entries to be measurements rather than adjectives. The 57 records hold 387 such entries between them, a median of seven per record. Third, there can remain unsettled issues. Here 1 of the 57 records stands as proposed rather than accepted. We report this rather than only the accepted subset, because a log that recorded solely the decisions already taken would describe a project that had stopped moving, and the open record is where active disagreements are noted.

#### The catalogue of failure modes

The second document, `how this project breaks.md`, catalogues every substantive defect found in the project, 21 at the time of writing. Its value lies less in the individual entries than in what they have in common, and the catalogue sorts them into four recurring disguises (Figure S3B). In each, the code selected a value by where it sat or what it was called rather than by what defined it, so the wrong value was as well-formed and as plausible as the right one. None of these defects raises an exception, and none produces an implausible figure.

The catalogue also records how each defect was actually found. Analysis of the different reasons underlying the defects is informative (Figure S3C). Only three of 21 were caught by a guard; the ratio did not improve as the catalogue grew. The catalogue’s own conclusion is that this ratio argues for writing guards rather than for additional auditing, and it states the design rule that follows: name what passes, never what fails, because a list of permitted values refuses a value nobody anticipated, whereas a list of forbidden values admits it. The sharpest instance is a flag computed as “verdict not in (underpowered, ungated, fail)”. When the control later returned a verdict nobody had anticipated, weak, two approaches silently began certifying their own rankings as validated.

**Fig. S3.**
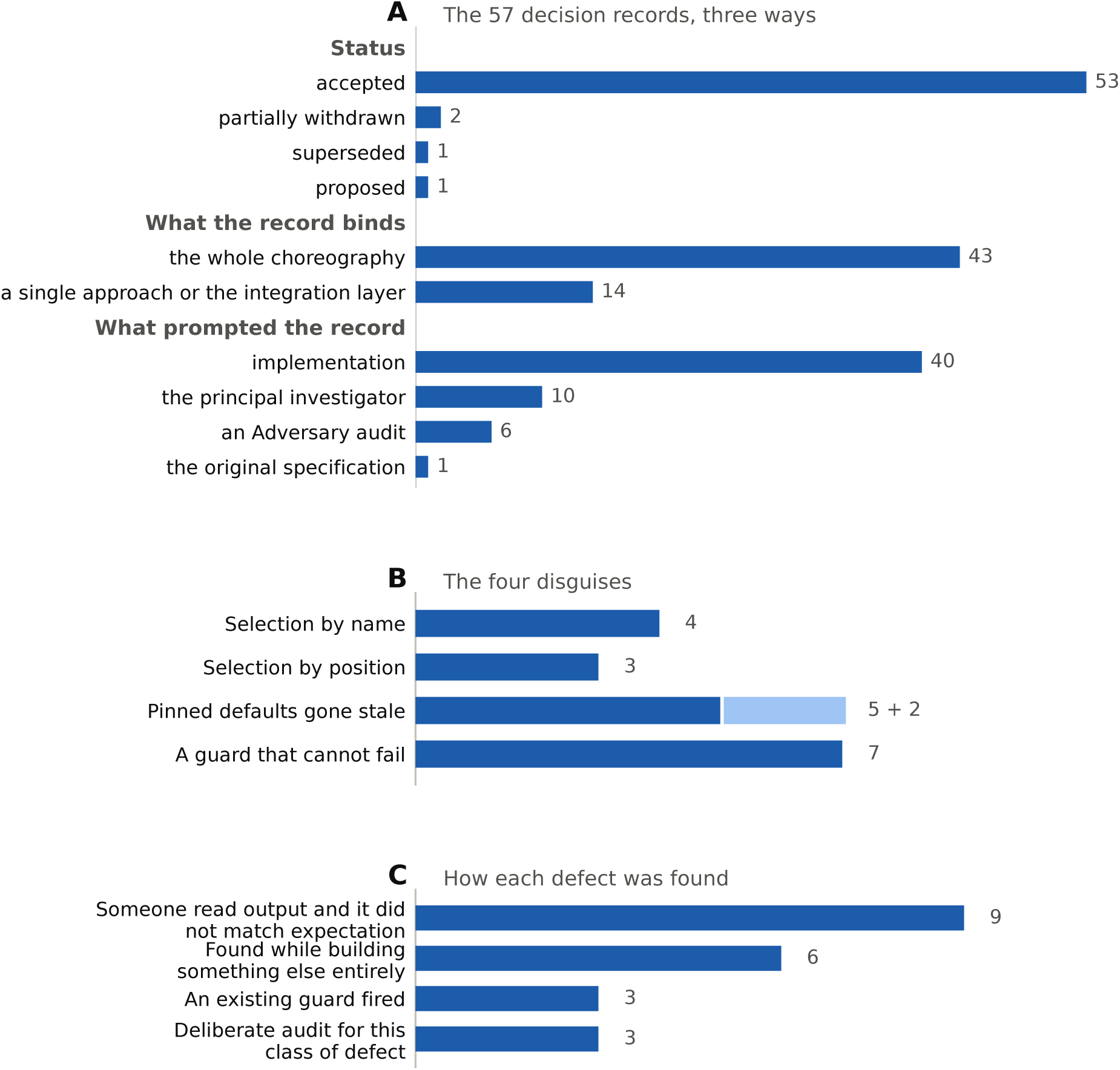
Summary of the written substrate of the *Pin1* choreography. **A.** The 57 decision records, one numbered file per binding choice under `decisions/`. Records that bind the whole choreography outnumber those scoped to a single approach by more than three to one. *What prompted the record* counts what raised the question, not who settled it: the PI prompted 10 records and signed off 51, and the two are different measurements of a supervisor’s role. The headers also carry 387 separate pieces of evidence, a median of seven per record; that is a count of a different thing and is not plotted here. **B.** The four recurring shapes behind the 21 defects catalogued in how this project breaks.md. The lighter segment is the two cache-key entries the catalogue treats as pins on their inputs; with those counted, every entry is assigned a shape. The largest group is the last, in which a guard was present throughout and simply could not fail. **C.** How each of those defects was found. Only three were caught by a guard, which is an argument about where guards are still missing rather than about how well the existing ones work.

#### Run manifests

The third document is written by the software rather than by a person. Every stage writes a manifest into the append-only tree beside its outputs, and it is what makes two results diagnosably different: without it, a change in the code, a change in the configuration and a change in the input are indistinguishable after the fact. A complete manifest is reproduced below. Note that it records whether the working tree carried uncommitted changes when the stage ran, which is the difference between a result that can be reproduced from a commit and one that cannot.

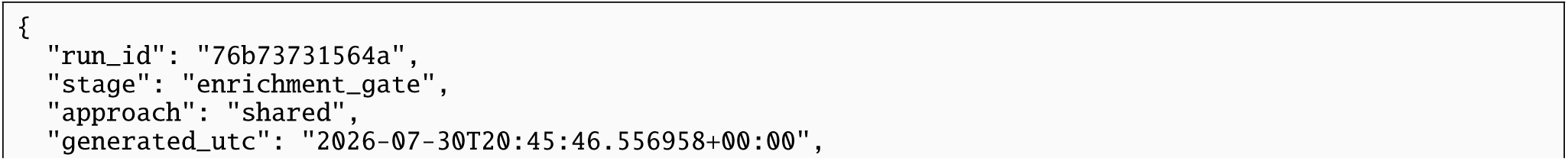
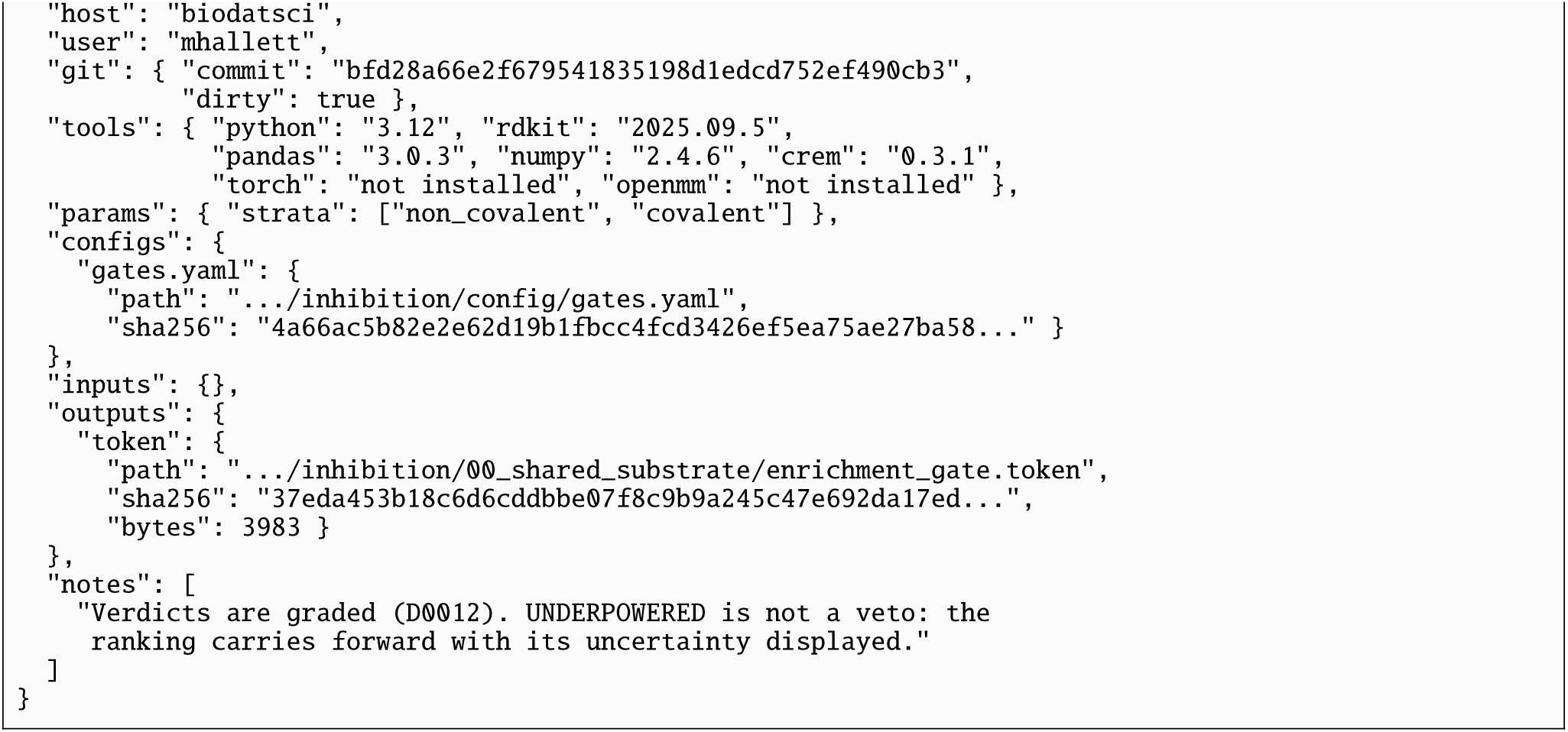

Taken together these three documents record what a source tree and a commit history do not. A test suite records what is checked; the catalogue records what was not checked and why the resulting numbers looked correct. Git records what the code became; the decision records say why, on whose authority, and what would have to change for the choice to be revisited. The manifests make the difference between two runs attributable to a specific input rather than to the passage of time.

### A.5 The course slides produced by the Conscience

The six slides described in §A.2 are reproduced on the following pages as the Conscience produced them, together with the speaker notes it wrote for the instructor and its statement of the sources behind each claim.

## Biochem 4450A: writing, equity and sex- and gender-based analysis

Six draft slides for you to adapt: three on scientific writing, two on a worked example, one on where disease names come from. Speaker notes sit under each slide, and each slide ends with where its claims come from. The styling is plain on purpose, so the content lifts easily into your own template.

## SLIDE 1 OF 6

### Sex, gender and race in a methods section

#### Sex and gender are two different measurements

- **Sex** is biological: chromosomes, hormones, anatomy.
- **Gender** is social: identity, roles, expression.
- Neither has only two values. Neither is fixed for life.

A good methods section answers three questions in one clause: which of the two you measured, how you got the value, and why it is in the analysis.

Don’t write this

"Gender was recorded for all participants." The values came from a hospital record, so they are neither gender nor cleanly measured sex.

*Made-up wording, not a quotation from anyone*.

Write this instead

"Sex was taken from the medical record and is reported as recorded there. Gender identity was not collected. Results are therefore not interpretable as gender effects."

#### Race is not a mechanism

Race is a social category with real health consequences. It is not a biological variable, and writing it as one puts a label where the mechanism you were meant to find should be.

Don’t write this

"Response was lower in Black patients, and the difference was statistically significant." The reader cannot tell whether the group label stands in for genetic ancestry, exposure, treatment delay, insurance, or nothing at all.

*Made-up wording, not a quotation from anyone*.

Write this instead

Name the thing you actually mean. If the claim is genetic, use measured genetic ancestry and say how you measured it. If the claim is about access or exposure, say that, and use the variable that carries it. If you do not know which it is, write that sentence: "we cannot distinguish these explanations with these data."

## SPEAKER NOTES

Lead with the accuracy argument, not the fairness argument. Students accept "this sentence does not say what you think it says" far more readily than "this sentence is insensitive", and in both examples above the accuracy argument is the stronger one anyway.

The second example is where the room may push back, because reporting group differences feels like reporting data. The reply is that a group label is a placeholder for an untested explanation. Ask the class to list what a race term could be standing in for, and you will get five candidate mechanisms in thirty seconds. That list is the point of the slide.

If you want a single line to close on: "a variable you did not measure cannot explain your result, and a label is not a measurement."

*Sources. From the centre’s equity resource list, sex and gender section: the paper at doi 10.1007/s10508-025-03331-y on moving away from a binary view of sex and gender and what follows from not doing so, plus 10.2471/BLT.22.289310 and 10.1016/j.ejim.2025.01.030. The three-question test above is that section’s own guidance. For race and biology, the nearest material on the list sits in its decolonizing section: entries on colonialism in the biosciences, and the United States National Human Genome Research Institute sessions on eugenics and scientific racism. Those are American, so treat them as material to learn from rather than a rule that binds you here. Two reporting standards are worth naming out loud. For sex and gender: SAGER, the Sex and Gender Equity in Research guidelines, Heidari and colleagues (2016) in Research Integrity and Peer Review 1:2, maintained by the European Association of Science Editors. For race and ethnicity: the AMA Manual of Style Committee’s updated guidance, Flanagin, Frey and Christiansen (2021) in JAMA 326(7):621 to 627, which states that race and ethnicity categories "have no scientific or biological meaning" and asks authors to say how the category was arrived at and to list categories alphabetically rather than by size. That second one is the authority behind this half of the slide*.

## SLIDE 2 OF 6

### Claiming more than your sample can support

The claim in a paper has to fit the sample that produced it. This is the most common failure in molecular oncology writing and the easiest to fix, because the fix is one sentence in the discussion.

Don’t write this

"These results establish PKX as a general determinant of chemosensitivity in breast cancer." Eight cell lines. One hospital. One country.

Write this instead

"These results establish PKX as a determinant of chemosensitivity in the eight luminal cell lines tested and in a single-centre cohort of 140 patients. Whether this extends to other subtypes, to other ancestry groups, or to other care settings is untested."

### The wording to watch for

"Generalisable to the population" is the phrase that should stop you. So should silence: a discussion section with no limitations paragraph is making a general claim by leaving one out.

### Where this matters most in this course: Weeks 2 and 3

Deciding whether a mutation matters means asking how common it is in "the population". That answer comes out of a reference database, and those databases were not built from everyone equally. Whichever ancestry group is under-sampled, a harmless variant that is common in that group can look rare and **be called pathogenic**, and a genuinely harmful one can be dismissed as noise. How often you get it wrong depends on the patient’s ancestry. That is a property of your method, not of the patient.

#### Two words, kept apart

*Pathogenic* is a judgement about a variant a person inherited, and it is the one that reference databases and frequency filters govern. *Driver* is a judgement about a mutation that arose in a tumour, and it comes out of a different analysis altogether. Use the wrong word and you have named the wrong analysis.

#### Two numbers, with their status attached

- **Peer reviewed:** 53% of high-impact coding variants found in Middle Eastern populations are absent from gnomAD, the reference database behind most clinical variant interpretation.
- **Not yet peer reviewed, so say so if you use it:** across nine established breast cancer risk genes including BRCA1, BRCA2, PALB2 and ATM, 43.4% of protein-truncating variants found only in South Asian populations are absent from ClinVar, against 20 to 30% for other ancestries.

**Two honest limits, and saying them makes you look better, not worse.** First, you cannot demand an analysis split by sex or by ancestry from data where the variable was never measured. The finding there is that the variable is unmeasured, and the fix belongs to the next study’s design. Second, when you are discussing somebody else’s earlier work, the criticism belongs on how you describe and interpret it, not on a demand that they re-run it.

## SPEAKER NOTES

The paired examples are the whole slide. Read the first aloud, then the second, and let the class notice that the second is more informative. **Say the real number: the first is 13 words and the second is 41, so the honest version takes 28 more words, about three times the length.** It is the easiest thing in the set for a student to count, so count it yourself first. The trade is still obviously worth making.

For the reference database point, the empirical source is the GWAS Diversity Monitor, which reports the live ancestry make-up of genome-wide association studies. Two cautions. It covers studies of inherited variation, not tumour sequencing, so do not cite it for a claim about tumour cohorts. And the figures are live: look at it yourself on the morning of the lecture rather than quoting a number from a slide deck, mine included.

The box at the bottom is what stops this becoming a licence to dismiss papers. Students who learn the criticism without the limit write reviewer comments that get overruled.

*Sources. From the centre’s equity resource list, decolonizing section: the GWAS Diversity Monitor, which the list names as the empirical citation for ancestry sampling bias in genomics. From the sex and gender section: both limits in the box, quoted almost directly. The gnomAD figure is from Middle Eastern Genetic Variation Improves Clinical Annotation of the Human Genome, peer reviewed and open access. The ClinVar figure is from a medRxiv preprint on breast cancer risk variants in South Asian populations; there is also a bioRxiv preprint analysing inherited variation from 9,899 patients across 33 TCGA cancer types if you want a cancer-specific version. Both preprints are unrefereed and the list says so. **One thing still missing from the list:** the ancestry make-up of the large tumour sequencing resources themselves, as distinct from the variant databases queried against them*.

## SLIDE 3 OF 6

### Who did what to whom, and describing a group by what it lacks

#### Who is the actor

The passive voice is fine where the actor genuinely does not matter: "the samples were centrifuged" is good writing. It becomes a problem the moment the actor is the interesting part.

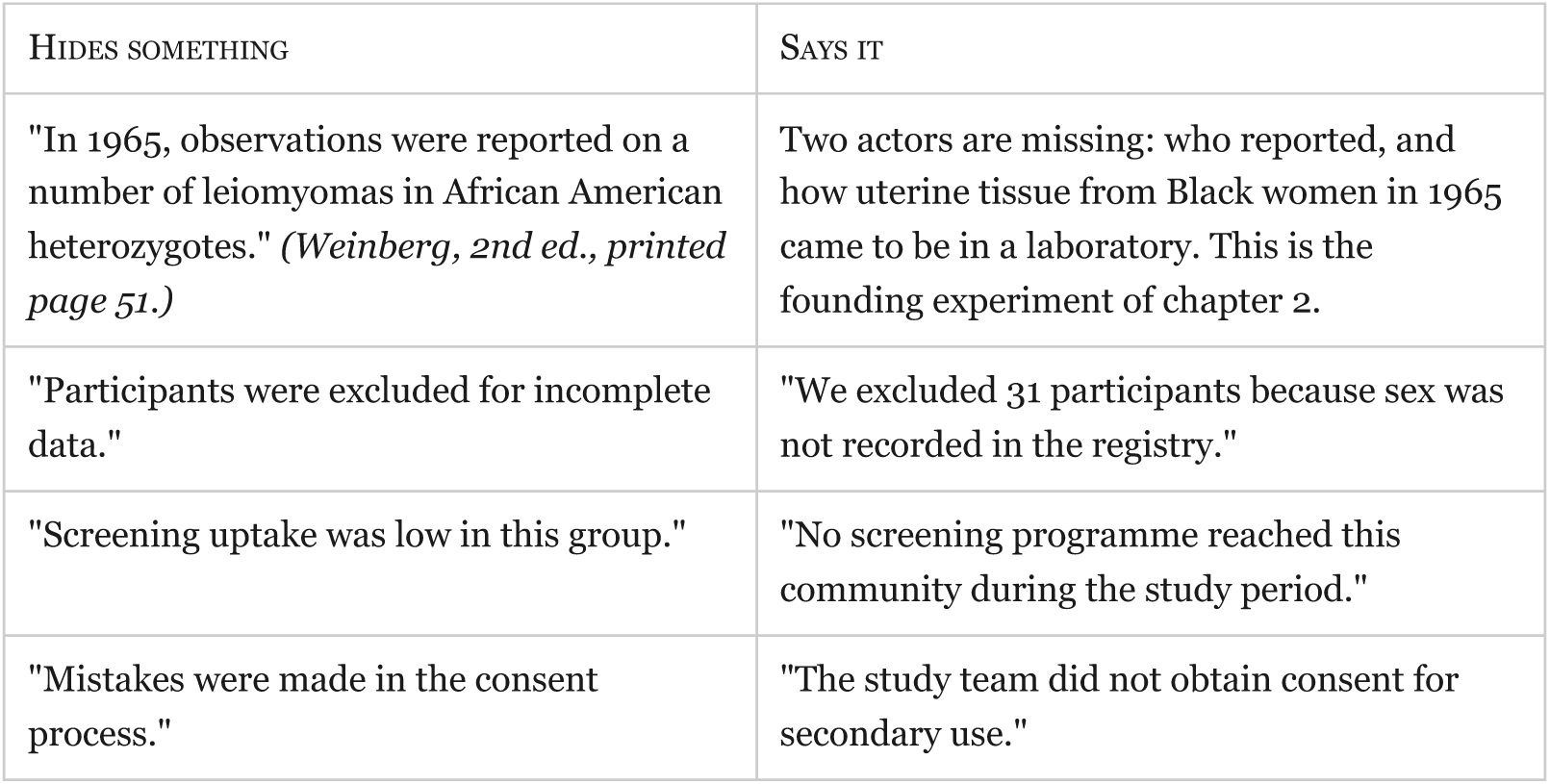

Each left-hand sentence is grammatical, publishable and less true. This matters here because the history of this field contains things that were done to people, and the passive voice is how they get recorded as things that happened.

#### Describing a group by what it lacks

Describing a group by what it is missing puts the cause inside the group and quietly ends the enquiry.

Don’t write this

"This population is hard to reach, has poor health literacy, and is non-compliant with follow-up."

Write this instead

"The nearest oncology centre is a six-hour drive. Appointment letters were sent in English only. Two of the three community clinics closed during the study."

The second version names things you could change. The first names a group you could blame. Same data.

#### One more: cells do not have an agenda

**Weinberg, 2nd edition, section 2.1, printed page 32.** Tumour cells "no longer obey the rules". Then: "**When portrayed in this way**, the renegade cells that form a tumor are the result of normal development gone awry… cancer cells somehow learn to thrive. **Normal cells are carefully programmed to collaborate** with one another… Cancer cells have a quite different and more focused agenda. They appear to be motivated by only one consideration."

**Five words there give the cell intentions: obey, renegade, learn, agenda, motivated.** ("Safeguards" appears in the book too, but attached to the organism rather than to the cell, so it does not belong in this list.)

**Read the quotation with both of its qualifiers in, because that is the honest reading.** "When portrayed in this way" is the book flagging its own picture as a picture, in the same sentence that uses it. And "carefully programmed to collaborate" does the same thing to normal cells, so the passage is an even-handed contrast rather than a one-sided account of cancer cells as outlaws. Dropping either qualifier is exactly the habit this slide teaches you to catch.

**Printed page 68, the chapter’s own key concepts.** "…tumors are not foreign bodies but growths derived from normal tissues." Printed page 69 sets that as thought question 3. And section 2.1’s own heading, on page 32, already says it: "Tumors arise from normal tissues."

The key concept is true and the prose is a habit. **That is the whole point, and it is about your writing, not about the textbook being wrong.** In your own writing, every word that gives a cell intentions is a place where a pathway and a signal should be. "The immune system fights the tumour" hides the entire content of this course, and that sentence is my own made-up example rather than a quotation from anybody. Keep "invasion" and "invasive carcinoma": there they are defined technical terms.

**Separately, about the literature and not about this book.** War and battle language is becoming more common in published science, and there is experimental evidence it lowers how credible readers judge the work to be. Say it that way round, because chapters 1 to 4 of your own textbook contain no "war on cancer", no "battle" and no "arsenal". They do contain "defenses" on page 58, so do not claim the book is free of that language altogether.

## SPEAKER NOTES

The table does the work. If you are short of time, drop the section on describing a group by what it lacks and keep rows one and three, which pair naturally: one is about the past of this field, the other about the present.

Row one is their own textbook, which is why it works. Do not assert anything about how that tissue was obtained. The point of the slide is that the sentence does not tell you. (HeLa would be the obvious choice here, but it does not appear in chapters 1 to 4; page 51 is both in the book and on topic, because a tumour arising from a single cell is a key concept of chapter 2.)

On the language that gives cells intentions, expect one student to say it is harmless convention. It is a fair challenge and the honest answer has two parts. The precision argument stands on its own: ask them to rewrite "cancer cells have a more focused agenda" with an actual pathway and they will see what the metaphor was covering. And the book agrees with them on page 68, which is the part that lands, because you are not asking them to take your word over their textbook’s.

### Do not stage this as the book contradicting itself

Page 32 carries the correct position itself, two paragraphs below the quotation, under a section heading that states it. What is happening is ordinary expository writing: a chapter opens with a rhetorical picture, hedges it, then corrects it. A student reading the page sees that, and if you have claimed a contradiction you have handed them the win.

### If a student asks how much war language is really in the book, here is the full answer

Six terms come back at zero across the 130 printed pages of chapters 1 to 4: "war on cancer", "battle", "arsenal", "magic bullet", "combat" and "immune surveillance". Six terms at zero is a statement about six terms, and no more than that. The book does carry other wording of the same kind, and page 58 is the counterexample a student would find first: "the body must be able to mount highly effective **defenses** that usually succeed in holding off the disease". Also "attack and corrupt" on page 57, "rampant, predatory and ungovernable" in the Rous epigraph on page 31, "attacked their DNA" on page 116, and "the powers of the immune system" as a sidebar title on page 83. Say exactly that, and never say there is no war language at all.

*Sources. From the centre’s equity resource list, inclusive language section: Almendingen (2025) in Cancer Control, open access, on what war language about cancer costs, and Mohapatra, Lydon-Staley and Bassett (2026), a study of 21.4 million papers from 2010 to 2025 finding this language rising and, in an experiment, lowering how credible readers judge the work. That 21.4 million figure comes from the resource list entry, not from my own memory, and **the second source is an arXiv preprint, not yet peer reviewed**, which you should say if you use the credibility finding. Also Western’s own Inclusive Language Guide, which is a Western instrument and so binds you here: **its section on socioeconomic status flags exactly the pattern this slide teaches**, giving "the poor" and "low-class people" against "people whose incomes are below the federal poverty threshold". That is about how you describe an individual’s income rather than how you describe a population in a methods section, so it is close but not exact, and it is Western’s own rule rather than one binding a collaborator elsewhere. Also the American Psychological Association’s Inclusive Language Guide, 2nd edition (2023), which the list marks as American material to learn from, and which states that neither person-first nor identity-first language is correct by default: the choice belongs to the people being described. **Still missing from the list:** anything on the passive voice in scientific writing, and anything on describing a group by what it lacks beyond the income wording in the Western guide above. Those two sections rest on the list’s remit for inclusive language and on my own judgement*.

## SLIDE 4 OF 6

### A worked example: the study, and what it left out

**This is a made-up example, not a real paper.** It was assembled from failure patterns that recur across the literature, and it is presented that way on purpose. No paper, author, journal, institution or country is being described, and none should be guessed at. The point is the principle, and an invented example presented as a real one would be worse than no example at all.

**One of these five omissions has been counted in real papers, so this is not just my say-so.** A systematic review of **228 cardiovascular studies using cultured cells**, published across **16 journals in 2018**, found the sex of the cells reported in **38.6%** of them, up from 19.8% in a comparable 2010 sample, with only **3 of the 16 journals** above half. Vallabhajosyula and colleagues, *The FASEB Journal*, 2020.

**Read the scope out loud, because this slide is about not claiming more than your sample supports.** That is cardiovascular research, cultured cells, 228 studies, 16 journals, one year. It is **not** a cancer figure and it covers **one** of the five omissions below. Presenting it as an oncology statistic would commit the exact error slide 2 teaches against.

It is also an aggregate across many papers, with no individual author or paper named as an offender, which is why it can go on a slide as it stands. It backs up the sentence above about patterns that recur, but it cannot replace the made-up example, because the other four rows and the shape of the exercise, auditing one paper end to end, would be lost.

### The study

A molecular oncology paper reports that expression of a kinase, call it PKX, predicts which patients with a solid tumour respond to a targeted inhibitor. The work has three parts.

1. **Cell lines.** Eight established human lines. Sensitivity to the inhibitor tracks with PKX expression.
2. **Mouse xenografts.** Two of the lines are grown in immunodeficient mice. High-PKX tumours shrink; low-PKX tumours do not.
3. **Patient cohort.** 140 archived tumour samples from one academic hospital, with treatment response from the charts. High PKX is associated with response. A cut-off is proposed for clinical use.

### What the paper never says

| PART OF THE STUDY | WHAT IS ABSENT |
| --- | --- |
| Cell lines | The donor sex of any of the eight lines. It is not in the paper and, for several lines, not easily findable. |
| Mice | The sex of the animals. The methods do not state it. The vendor's default for that strain was used. |
| Patient cohort | A demographic table headed "gender", with two values, taken from the hospital record. No statement of how it was arrived at and no mention of why it is there. |
| Patient cohort | Any description of the cohort's ancestry, language, or distance from the hospital. The single recruiting centre is named but the area it draws from is not described. |
| Discussion | A limitations paragraph. The proposed cut-off is stated without a boundary on who it applies to. |

## SPEAKER NOTES

Say clearly, twice if needed, that this example is made up. The first time students meet this kind of critique they want to know which paper it was, and the answer being "it is every third paper" is more useful than a name. It also keeps the room thinking about design rather than about whether the authors were careless, which is not the lesson.

Put up the study first, with the omission table covered, and ask the class what is missing. In my experience of this exercise the room reliably finds the single-centre cohort and reliably misses the sex of the cell lines and the mice, which is the useful half of the result: they have been trained to audit the clinical part and not the bench part.

Worth saying out loud: none of these omissions is misconduct and none of them would stop this paper being published. That is the problem being described.

*Sources. The example is built to show the failures the centre’s equity resource list names in its sex and gender section: using sex and gender interchangeably, reporting one having measured the other, collecting either as a single two-value field with no stated reason, using either as a variable without saying which thing it stands for or how it was arrived at, and silently excluding groups rather than making that a stated design decision. The sampling and over-claiming half comes from the same list’s decolonizing section. The audit in the second box is Vallabhajosyula, Ponamgi, Shrivastava, Sundaragiri and Miller (2020), Reporting of sex as a variable in cardiovascular studies using cultured cells, The FASEB Journal 34. It covers one of the five rows above, in one field*.

## SLIDE 5 OF 6

### The same example: what followed, and how to fix it

**Still the made-up example, still not a real paper.** The findings below are constructed from failure patterns, not taken from any publication. No paper, author, journal, institution or country is being described. This label is on the slide itself rather than in the speaker notes because a student who photographs this slide, or who arrives late, would otherwise take these for a real audit. For the one of them that has been counted in real papers, see slide 4.

#### What followed from the omissions

- **A sex-dependent effect could not be seen, in either direction.** With the donor sex of the lines unknown and the animals all one unstated sex, the study has no capacity to detect one. The result is not that there is no sex difference. The result is that nobody can tell, including the authors. *This is what the 38.6% on slide 4 is measuring: how often sex was reported, not how often somebody got the answer wrong. The other 61.4% of those studies did not get the wrong answer. Nobody can check them*.
- **The demographic variable answers no question.** "Gender" from a chart is not gender and is not clean sex data. It occupies a row in the table and supports no inference.
- **The proposed cut-off travels further than the data.** A threshold from one hospital’s archive, with no description of who that archive represents, gets cited as a general threshold. The next group applies it to a different population and the performance drops, and nobody can work out why, because the original cohort was never described.
- **The failure is invisible in everything published afterwards.** Nothing in the paper flags the limits, so later reviews repeat the claim at full strength.

#### The corrected design, in five lines

1. **Report the donor sex of every cell line**, and say so where it is unknown. One extra column in a supplementary table.
2. **State the sex of the animals, and use both** where the number of animals allows. Where it does not, state the sex used and say the study cannot address sex dependence. That sentence costs nothing and protects the claim.
3. **Say which thing you measured in the human cohort.** "Sex as recorded in the medical record" or "self-reported gender identity, collected at enrolment". Say how you got the value and why it is in the analysis. If intersex, trans or non-binary participants were combined with others or excluded, say so and say why: an undeclared exclusion is a methods problem, a declared one is a design decision.
4. **Describe the cohort.** Recruiting centre, the area it draws from, and the variables that bear on the claim. Where the claim is genetic, use measured genetic ancestry. Where the claim is about access or care, use the variable that carries that, and do not let one stand in for the other.
5. **Bound the claim in the discussion.** "Validated in a single-centre cohort; performance in other settings and populations is untested." **Thirteen words**, and it is the difference between a finding and an overclaim.

#### The line to leave them with

##### Four of these five cost you a sentence each

Saying which thing you measured, describing the cohort, bounding the claim, and saying where a cell line’s donor sex is unknown are all writing, and the last of those is the honest move precisely when the fact cannot be found.

##### The fifth is a real design decision and it does cost something

Using both sexes costs animals, money and statistical power, which is why correction 2 says "where the number of animals allows" and then tells you what to write when it does not. Say that plainly. The argument gets stronger when it stops overclaiming.

Sex- and gender-based analysis at the bench, which the Canadian granting councils call SGBA+, is mostly the discipline of writing down what you already know and declining to claim what you did not measure.

## SPEAKER NOTES

The most important sentence on this slide is "the result is not that there is no sex difference, the result is that nobody can tell". Students routinely read a silent methods section as a null result. That single confusion is worth the whole lecture.

Point three is where you connect back to the funder requirements if you covered them. The Canadian granting councils expect equity, diversity and inclusion to be addressed in research, and the natural sciences council and the New Frontiers in Research Fund both publish guidance on it. Students heading to graduate school will write this section of a grant application within two years.

Close on the cost, and be accurate about it. Four sentences and one supplementary column is most of the work, and using both sexes is the one item that needs a design decision. If the room takes away that four fifths of this is free and the fifth is a choice you should make deliberately, the lecture has worked.

*Sources. From the centre’s equity resource list, sex and gender section: the two-variable framing, which is the substance of corrections one to three, and the funder guidance named in the notes. Specifically, the Natural Sciences and Engineering Research Council’s guide on integrating equity, diversity and inclusion considerations in research, the New Frontiers in Research Fund best practices page, and **the Canadian Institutes of Health Research’s requirement that every Project Grant application describe how sex and gender are built into design, data collection, analysis and dissemination, with a "not applicable" answer justified rather than left blank**. All Canadian policy, so it binds an applicant here. Corrections four and five come from the list’s decolonizing section. The reporting standard to end on is SAGER, the Sex and Gender Equity in Research guidelines, Heidari and colleagues (2016), maintained by the European Association of Science Editors. General authority, not specific to Canada*.

## SLIDE 6 OF 6

### Whose name is on the disease?

Chapters 1 to 4 of your textbook run on names of people. The **Barr body** (pages 10 and 51), **Hodgkin’s and non-Hodgkin’s lymphoma** (page 39), **Barrett’s esophagus** (page 46), **Burkitt’s lymphoma** (page 81), **Wilms tumour** and **Ewing’s sarcoma** (page 13), and the **Kaplan-Meier** plot (page 110). You will say all seven this term.

Every one of them names a disease or a method after the person credited with describing it first. Looking at where that credit actually came from tells you something about how science assigns credit generally. **It is not one story. It is at least three, and the useful skill is telling them apart.**

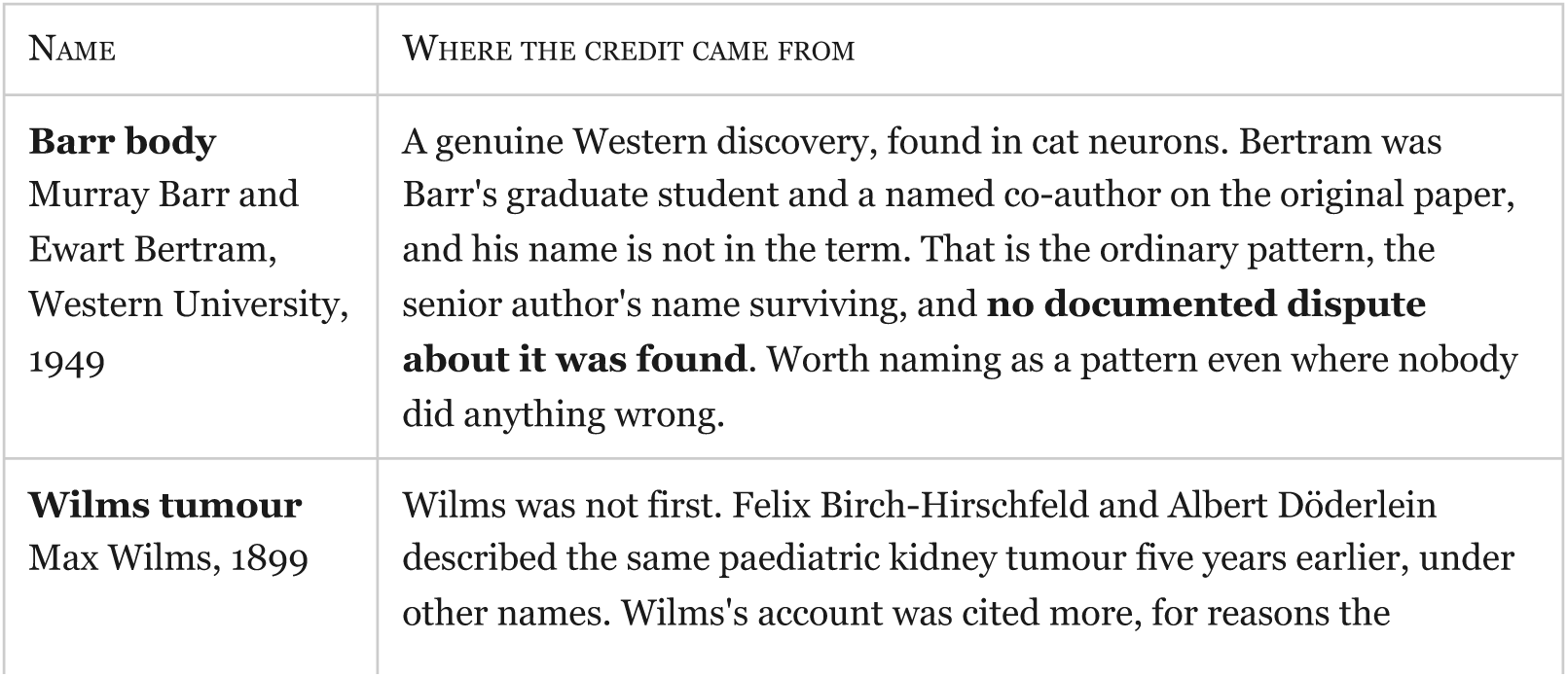
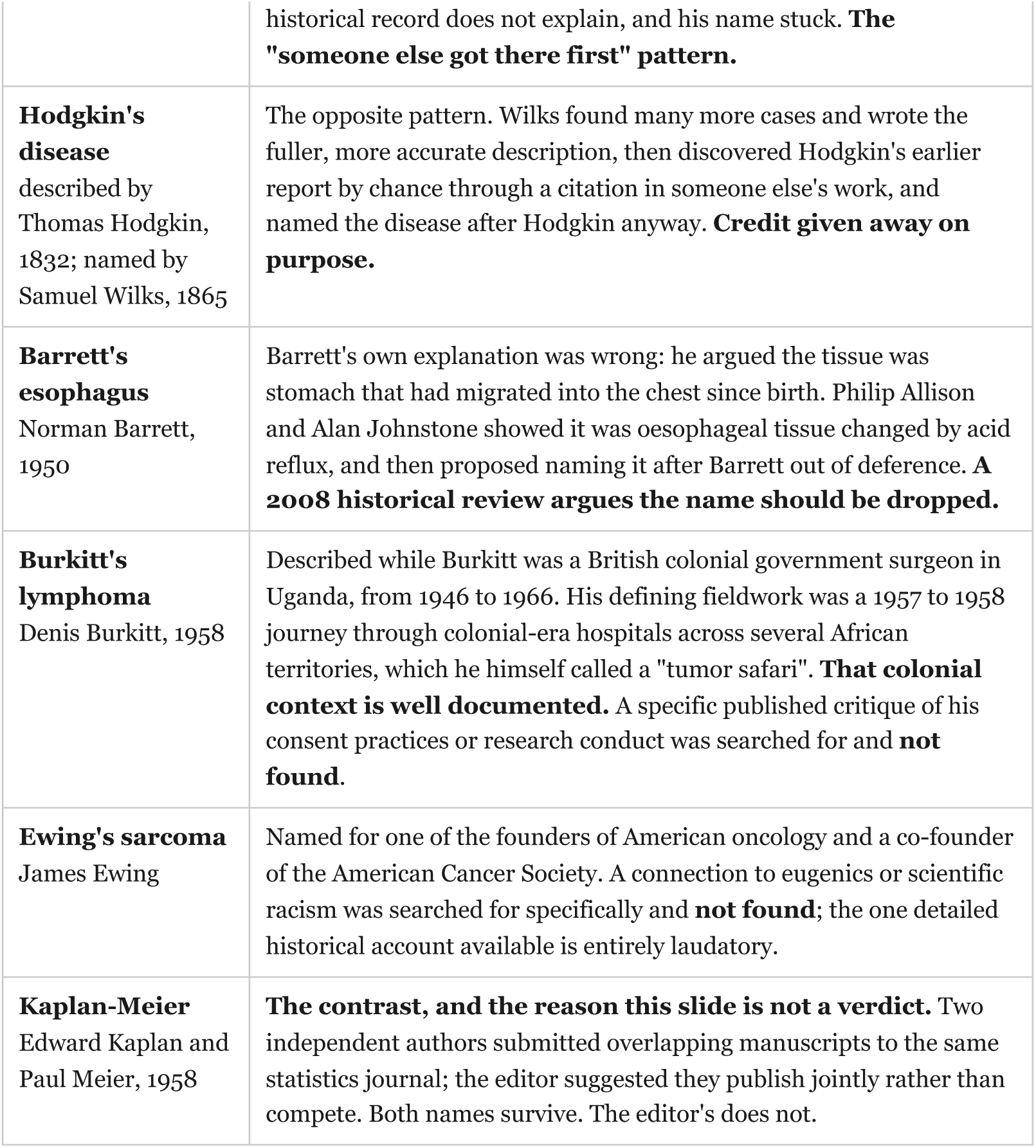

**Two honest absences, and they are the point of saying them out loud.** For the Barr body and for Ewing’s sarcoma, somebody searched specifically for a dispute or an ethical problem and found none. **Nobody found anything is not the same as there is nothing to find**, and that difference is the same one slide 5 makes about unmeasured variables. Do not upgrade either one in the telling.

#### The question, not the answer

Some people argue these names should be replaced with descriptive terms, because they are inconsistent across countries and journals and often describe nothing about the condition (Fargen and Hoh, 2014). Others argue the specific case of colonial and racist names in science needs retiring on ethical grounds regardless of convenience (Chala and colleagues, 2024, *PNAS*, open access). Nobody argues Kaplan-Meier is a problem. **Ask the room which of the seven above belongs in which argument.**

## SPEAKER NOTES

Lead with Barr. It is a Western discovery, it is in their chapter 1 and chapter 2 reading, and it is the safest of the seven because nothing scandalous is attached to it. The point lands better from a case where nobody did anything wrong: a graduate student co-discovered the thing and the name records one person.

Then run Wilms, Hodgkin and Barrett together, because they are three different shapes of the same question and the class can sort them in a minute. Finish on Kaplan-Meier, which is in the figure you showed alongside slide 4 in the insertions document, and which is the reason this is a question rather than a verdict.

### Two things to say precisely

Two of the sources behind this slide are paywalled and were read as abstracts only, and one background source is a physician-run reference site that the centre’s resource list marks as background reading rather than an authority. Say "the historical accounts I could read say" rather than "it is established that". And for Burkitt and Ewing, say "no published criticism was found", not "there was none".

### What I could not get, and it would have been the best item on this slide

Epstein-Barr virus appears in your own chapter 4 teaching block more often than any name above, on pages 81, 102, 120, 121 and 127. The book never gives a first name or an initial for either person. Who they were is not in the centre’s source list, and I will not assert a person’s identity from my own memory on a slide about people being written out of names. The centre’s literature search has been asked for it. If it comes back sourced, it belongs here.

*Sources, and the licence position, which matters for this slide in particular. From the centre’s equity resource list, inclusive language section: Zantinga and Coppes (1992) in Medical and Pediatric Oncology 20(6) for Wilms, abstract only; Bani-Hani and Bani-Hani (2008) in the Journal of Gastroenterology and Hepatology 23(5) for Barrett, abstract only; the PubMed Central records PMC1807800 and PMC4450782 for Hodgkin and Wilks, **cited by identifier because the list flags their author attributions as unconfirmed**; Stalpers and Kaplan (2018) in BSHM Bulletin 33(2) for Kaplan-Meier; PMC2995899 for Ewing; Fargen and Hoh (2014) in Clinical Anatomy 27(8) for the debate, abstract only; and Chala and colleagues (2024) in PNAS for the case against colonial and racist names. The LITFL medical eponym library was used as background, and the list marks it as background rather than an authority, so nothing here rests on it alone*.

### On the Inclusive Anatomy Eponym Directory, and why it is named here but not used

*The Directory is a Western-led project, founded and led by Dr. Charys Martin, pairing anatomical names with descriptive replacement terms. It covers **anatomical structures** across seven body regions, and **none of the seven names above appears on it**, because diseases and a statistical method are a different sort of name. That was confirmed by looking at its region index. So it is cited here for what it is, and nothing on this slide is adapted from it. **That is deliberate, and it settles a licence conflict rather than ignoring one:** the Directory is CC BY-NC-SA, which forbids putting further restrictions on adapted material, and this course’s own materials carry a no-share stamp. Writing these histories from the medical-history literature in the instructor’s own words, with the sources named, means the licence never comes into play. Do not paste Directory text or tables onto these slides*.

### How these slides fit the Fall 2026 course outline

Slides 1 to 3 are the outline’s own learning outcomes, not an addition to them. Outcome 1 asks that students "understand and evaluate news reports and new breakthroughs reported in the popular press in proper scientific context". Outcome 4 asks them to "clearly communicate technical concepts and experimental results to diverse audiences in writing, in a style consistent with practice in the field". Say that when you use these, because it tells students the material is examinable. Slide 2’s section on reference databases belongs in your Week 2 and 3 lectures on genome maintenance, where deciding whether a variant matters is the subject rather than an aside.

### The edition problem, confirmed from both documents

The textbook file on the machine is the **second** edition, Garland Science, copyright 2014, ISBN 978-0-8153-4219-9, and the outline’s section 10 assigns the **third**, from Norton. Every page number in this deck is second edition and will not match a student’s copy. Say so in the first lecture, give readings by topic, and ask students to tell you when a number does not match.

### No Canadian or Ontario cancer statistic appears on any slide, and here is why

There is one properly governed source available: *Cancer in First Nations People in Ontario*, co-produced by Chiefs of Ontario, Ontario Health’s Indigenous Cancer Care Unit and ICES, built under a data governance and sharing agreement implementing the First Nations principles of OCAP®. Its headline findings may be cited as that report’s findings, attributed to it by name. **But whether First Nations cancer data belongs in this course, whose data, and taught which way, is not a decision this review can make**, because it needs the people it describes; the route runs through Western’s Office of Indigenous Initiatives. The accompanying document `02_weinberg_ch1_4_edid_insertions.md`, point 5b, sets out what can be said honestly now, and the student handout carries the same wording. No Inuit-specific source was found, and that absence is recorded as a finding rather than passed over.

### Still missing

Your existing lecture slides, so I cannot tell you where these three writing slides overlap with what you already show.

## Notes

https://github.com/hallettmiket/murmurent

https://github.com/hallettmiket/murmurent_dev

https://github.com/hallettmiket/murmurent_public

### Competing Interest Statement

The authors have declared no competing interest.

